# Remotely controlled drug release in deep brain regions of non-human primates

**DOI:** 10.1101/2023.10.09.561539

**Authors:** Matthew G. Wilson, Taylor D. Webb, Henrik Odéen, Jan Kubanek

**Affiliations:** Department of Biomedical Engineering, University of Utah, 36 S Wasatch Dr, Salt Lake City, UT 84112, USA; Department of Radiology and Imaging Sciences, University of Utah, 729 Arapeen Drive, Salt Lake City, UT 84108, USA

**Keywords:** Drug release, ultrasound, nanoparticle carriers, pharmacomodulation, neuromodulation

## Abstract

Many areas of science and medicine would benefit from selective release of drugs in specific regions of interest. Nanoparticle drug carriers activated by focused ultrasound—remotely applied, depth-penetrating energy—may provide such selective interventions. Here, we developed stable, ultrasound-responsive nanoparticles that can be used to release drugs effectively and safely in non-human primates. The nanoparticles were used to release propofol in deep brain visual regions. The release reversibly modulated the subjects’ visual choice behavior and was specific to the targeted region and to the released drug. Gadolinium-enhanced MRI imaging suggested an intact blood-brain barrier. Blood draws showed normal clinical chemistry and hematology. In summary, this study provides a safe and effective approach to release drugs on demand in selected deep brain regions at levels sufficient to modulate behavior.

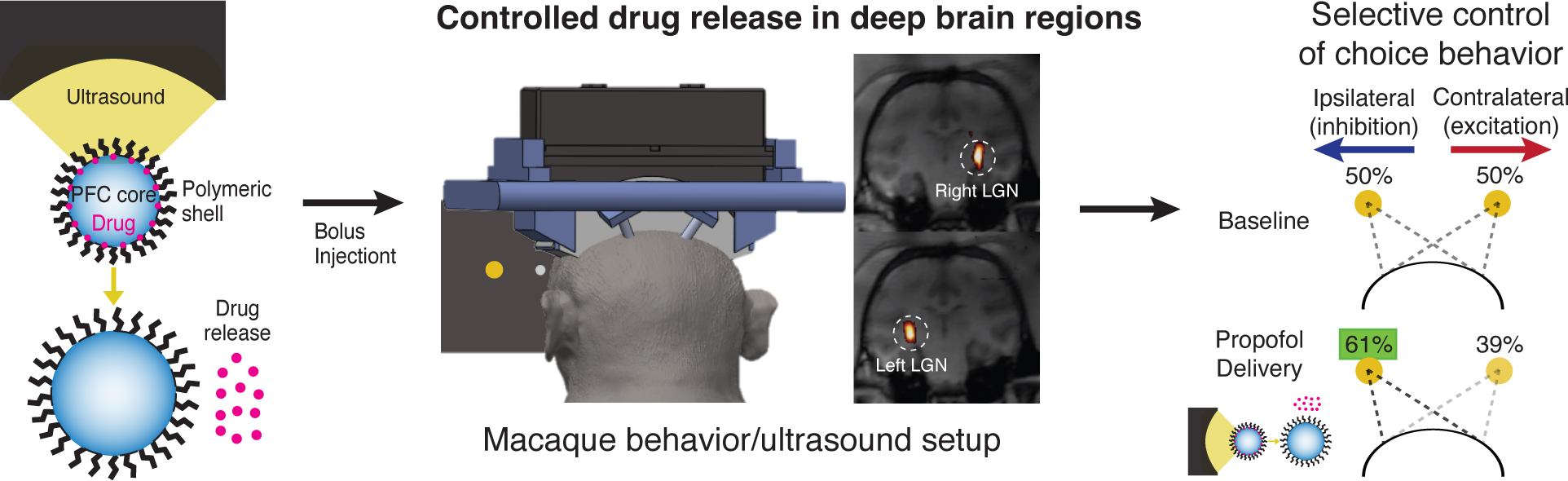

## Introduction

Systemic administrations of drugs often carry substantial side effects, which limits the dose that can be delivered, the range of drugs that can be administered safely, and, consequently, the spectrum of patients who can receive effective treatments. An ideal approach would release medication specifically in the target of interest and use remotely applied, noninvasive form of energy as the release trigger.

Ultrasound-sensitive nanoparticle carriers are emerging as a candidate for such a selective approach [1]. A key benefit of this approach is that ultrasound is applied remotely, outside of the body, while being able to penetrate into a selected organ at depth. Drug release from the nanoparticles occurs specifically at the ultrasound focus (Fig. 1A), which can comprise millimeter-sized volumes [1, 2].

**Figure 1.**
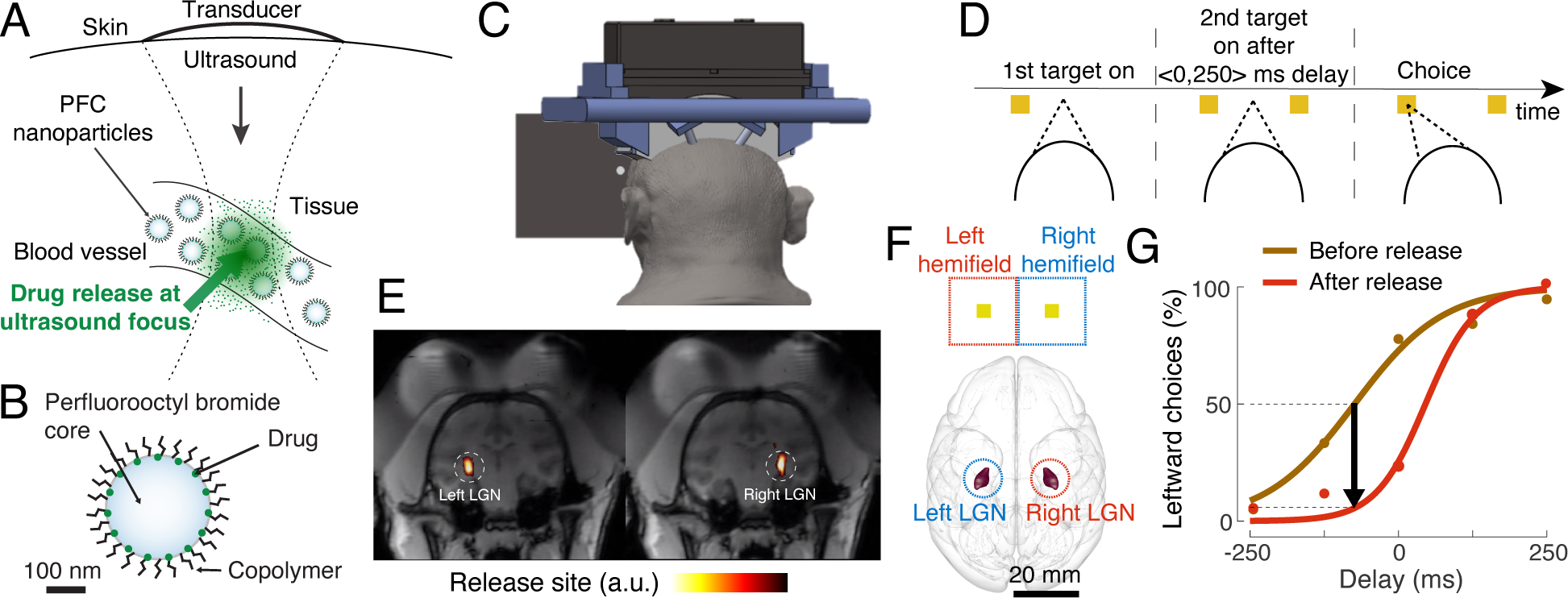
Ultrasound-triggered drug release from nanoparticle carriers in deep brain regions of non-human primates. A) Concept. Remotely applied focused ultrasound enables selective release of drugs from nanoparticle carriers specifically at its focus. B) Nanoparticle formulation. The nanoparticles consist of a perfluorocarbon (PFC) with a high boiling point— perfluorooctylbromide. Perfluorooctylbromide bestows the nanoparticles with high stability and biological safety [22–24]. The nanoparticle is further stabilized using a polyethylene glycol/polylactic acid co-polymer matrix. C) Ultrasound-controlled release in deep brain circuits of task-performing NHPs. A 256-element ultrasonic transducer array [27, 28] delivers ultrasound programmatically into deep brain regions of NHPs, enabling selective drug release is specified brain regions. The array is mounted into implanted head posts to ensure reproducible positioning of the transducer with respect to the head from session to session. D) Visual choice task. One target appears on the left and one target on the right part of the screen, with brief, controlled delay between the onsets. Subjects look at the target that appeared first. E) Validation of ultrasound targeting of the left and right lateral geniculate nucleus (LGN) using MRI thermometry. F) Brain hemisphere-specific representation. The left/right LGN relays visual information about the right/left visual hemifield into primary visual cortex. G) Example psychometric curve during a 3-minute baseline (brown) and a 3-minute period following the release of propofol (red) in the right LGN fitted to sigmoid curves. Henceforth, the choice bias following the release is quantified as the proportion of choices at the point of equal preference established during the baseline (black arrow).

Ultrasound-sensitive nanoparticles are commonly filled with a chemically inert perfluorocarbon core [3–11], which bestows them with sensitivity to ultrasound. Proof-of-concept release from such nanoparticles has been demonstrated in rodents [5, 7–10, 12]. However, it has been difficult to determine which combination of perfluorocarbon cores and ultrasound release parameters could mediate both effective and safe applications. This issue has, thus far, precluded translations of this approach to primates and humans.

This article finds that low-frequency ultrasound, in conjunction with stable, high-boiling-point perfluorocarbon nanoparticles, provides release that is both effective and safe. The approach is deployed in non-human primates for targeted neuromodulation of deep brain regions. We targeted deep brain regions with propofol to i) demonstrate the non-invasive nature of the approach and ii) realize its promise in selective treatments of mental and neurological disorders, which are commonly associated with malfunctioning deep brain circuits [13–19]. A demonstration of the safety and efficacy of this approach in awake, behaving primates, as opposed to previous studies that used rodents and primarily low-boiling point nanoparticles, provides a critical step toward clinical translation.

We loaded the nanoparticles with propofol as this drug inhibits neural circuits with rapid onset and offset [20, 21]. This provides a specific prediction of the expected effects. In addition, since propofol is readily used in clinics, its local delivery could open new diagnostic opportunities. A systematic inhibition of specific brain regions would provide direct information on the causal involvement of those regions in specific behaviors or behavioral disorders.

## Results

### Ultrasound-triggered drug release *in vitro*

We developed ultrasound-sensitive nanoparticle carriers (Fig. 1B) with a core comprising a high boiling point perfluorocarbon—perfluorooctylbromide (PFOB). PFOB has been used safely in large quantities in humans as a blood substitute [22–24], and its high boiling point (142*^◦^*C) contributes to the stability of the nanoparticles in the blood stream. The nanoparticles are further stabilized using a co-polymer shell (Fig. 1B).

We loaded the nanoparticles with the anesthetic propofol [8, 25]. First, we quantified the effectiveness of the propofol release *in-vitro*, using an approach described previously in which a drug is released into an organic solvent [26]. Ultrasound of increasing pressure increased the effectiveness of the release, which reached over 80% for pressures above amplitude of 1.3 MPa (Fig. S1A). The modulation of the release by the ultrasound pressure was significant (one-way ANOVA: *F* (6, 42) = 443.83*, p <* 0.0001).

We have observed no difference in the ultrasound responsiveness of the nanoparticles in plasma compared to PBS, as shown in Fig. S1A. This is confirmed using a two-way ANOVA with factors of ultrasound pressure and the dispersant; the dispersant factor was not significant (*F* (1, 42) = 0.20, *p* = 0.65). The amount of time the nanoparticles are in contact with plasma did not affect rates of drug release either with or without 1.5 MPa ultrasound (Fig. S2). The effect of drug release as quantified by a one-way ANOVA over time in plasma was not significant either with ultrasound (*F* (2, 9) = 0.47*, p* = 0.64) or without (*F* (2, 9) = 3.69*, p* = 0.068). This suggests that the particles remain stable when in contact with plasma. Additional details concerning the characterization of nanoparticles are available in Supplemental Information.

### Ultrasound-triggered drug release in deep brain regions of non-human primates

We next evaluated the release capacity of the approach *in-vivo* in non-human primates (NHPs), testing the ability to release drugs in specific deep brain regions through the intact skull and skin. To do so, we developed a system (Fig. 1C) that enables controlled and reproducible delivery of focused ultrasound into deep brain targets of awake NHPs [27, 28]. We engaged the subjects in an established choice task that is often used in neurology to evaluate the effects of stroke in visual regions [29, 30] and in neuroscience to quantify neuromodulatory effects [27, 28, 31–33]. In this task, subjects decide whether a left or a right target appeared first (Fig. 1D) and make an eye movement to that target. We specifically targeted the left and right lateral geniculate nuclei (LGN; Fig. 1E), which are the input nuclei of visual information to the brain.

Each LGN represents the contralateral visual hemifield and target (Fig. 1F), and so provides a well-defined framework for interpreting the polarity and magnitude of neuromodulatory effects on visual choice behavior [27, 28]. For instance, if propofol, which is an anesthetic, neuroinhibitory drug, is released in the right LGN, one can expect an impaired perception of the left target. This should lead to an ipsilateral, rightward preference in the visual choice task (Fig. 1D,F).

To test this prediction, we established a behavioral baseline (brown in Fig. 1G). We then injected the nanoparticles into the blood stream using a bolus such that the concentration of the encapsulated propofol was 0.5 mg/kg. Following a 1-minute delivery of pulsed ultrasound into the right LGN, we indeed found a strong ipsilateral bias in the animal’s choices (red in Fig. 1G). The animal chose the rightward target more frequently following the release of propofol in the right LGN, consistent with a release of a neuroinhibitory drug in the LGN.

The ultrasonic array enables selective drug release in specified brain regions, which allowed us to evaluate this effect systematically for both the right and the left LGNs. Across all recorded sessions, we found that the released propofol indeed induced an ipsilateral bias in the animals’ choices (Fig. 2, blue). This effect was specific to the propofol released from the nanoparticles; ultrasound alone of the same parameters had, if anything, the opposite effect (Fig. 2, red). Moreover, the effect was specific to the sonicated LGN side. We quantified these effects using an ANOVA that incorporated all 80 recorded sessions with factors of subject (Monkey 1 or 2), ultrasound pressure (1.2 or 1.5 MPa), drug (propofol nanoparticles or saline), and sonicated LGN side (left or right), and evaluated the choice behavior during the time window of expected propofol effects (2-5 minutes following the ultrasound onset [34]). The ANOVA detected a highly significant (*F* (1, 66) = 18.73*, p* = 5.2*e −* 05) interaction between the carrier factor and the sonicated LGN side (left or right LGN). This significant interaction demonstrates that the ultrasound released propofol selectively in each LGN and was capable of selectively modulating choice behavior. The discrepancy may be attributed to differences in targeting or estimation of ultrasound pressure delivered to the LGN.

**Figure 2.**
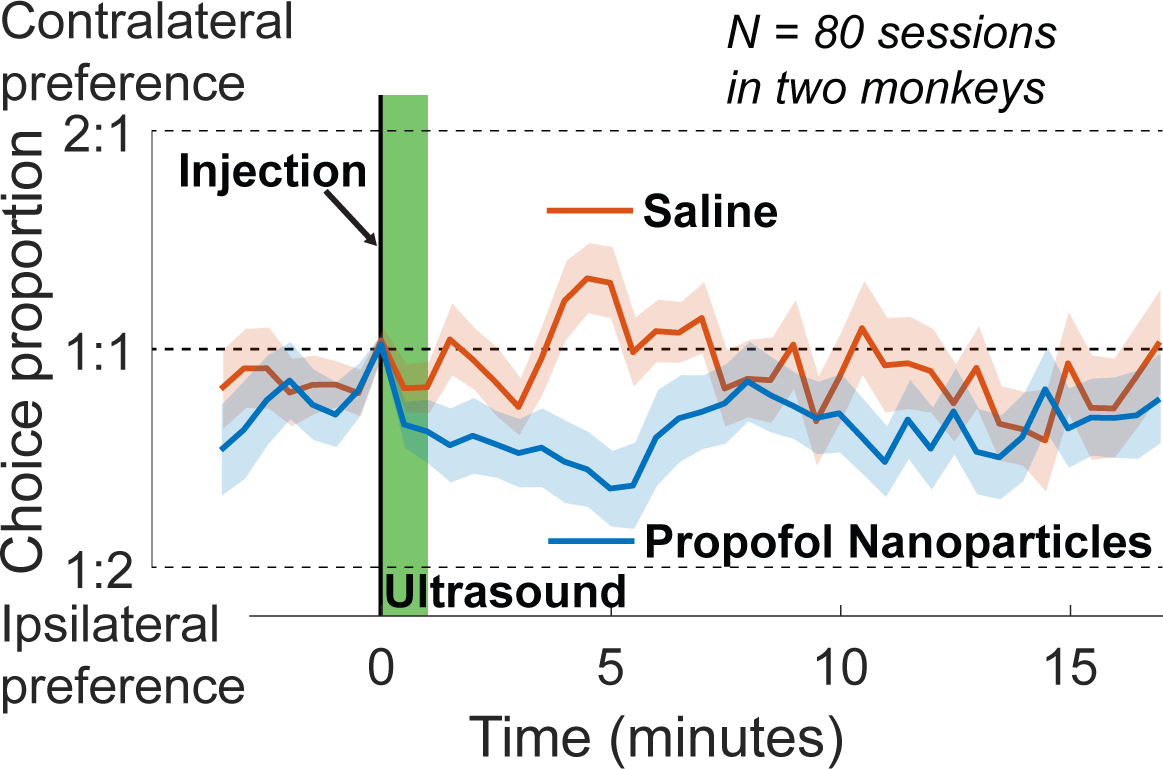
The release of propofol in deep brain regions of NHPs modulates choice behavior. Mean*±*s.e.m. proportion of choices contralateral to the targeted LGN as a function of time for the propofol-loaded nanoparticles (blue) and saline (red). The choice proportion was quantified in 3-minute moving-average windows, and was evaluated with respect to the point of equal preference obtained during a 3-minute baseline established prior to the injection of the nanoparticles (Fig. 1F). The injection and the ultrasound exposure times are indicated with a vertical black line and green bar, respectively.

The inhibitory effect is notable also when we separated the data into the right and left LGN release sessions, and was observed for both levels of tested ultrasound pressure (Fig. 3). A two-way ANOVA, evaluated in the same time window, again detected a significant interaction of the LGN side and carrier type (1.2 MPa: *F* (1, 31) = 8.65*, p* = 0.006; 1.5 MPa: *F* (1, 31) = 10.00*, p* = 0.003). These results confirm that ultrasound-triggered propofol delivery induced transient side-specific neuroinhibitory effects. Moreover, the effect shows the expected polarity given that propofol is a neuroinhibitory drug. Further, we have demonstrated that nanoparticles containing no propofol are not sufficient to induce any bias in behavior different from saline alone Fig. S4. A summary of the data used in each of these analyses is given in Fig. S5.

**Figure 3.**
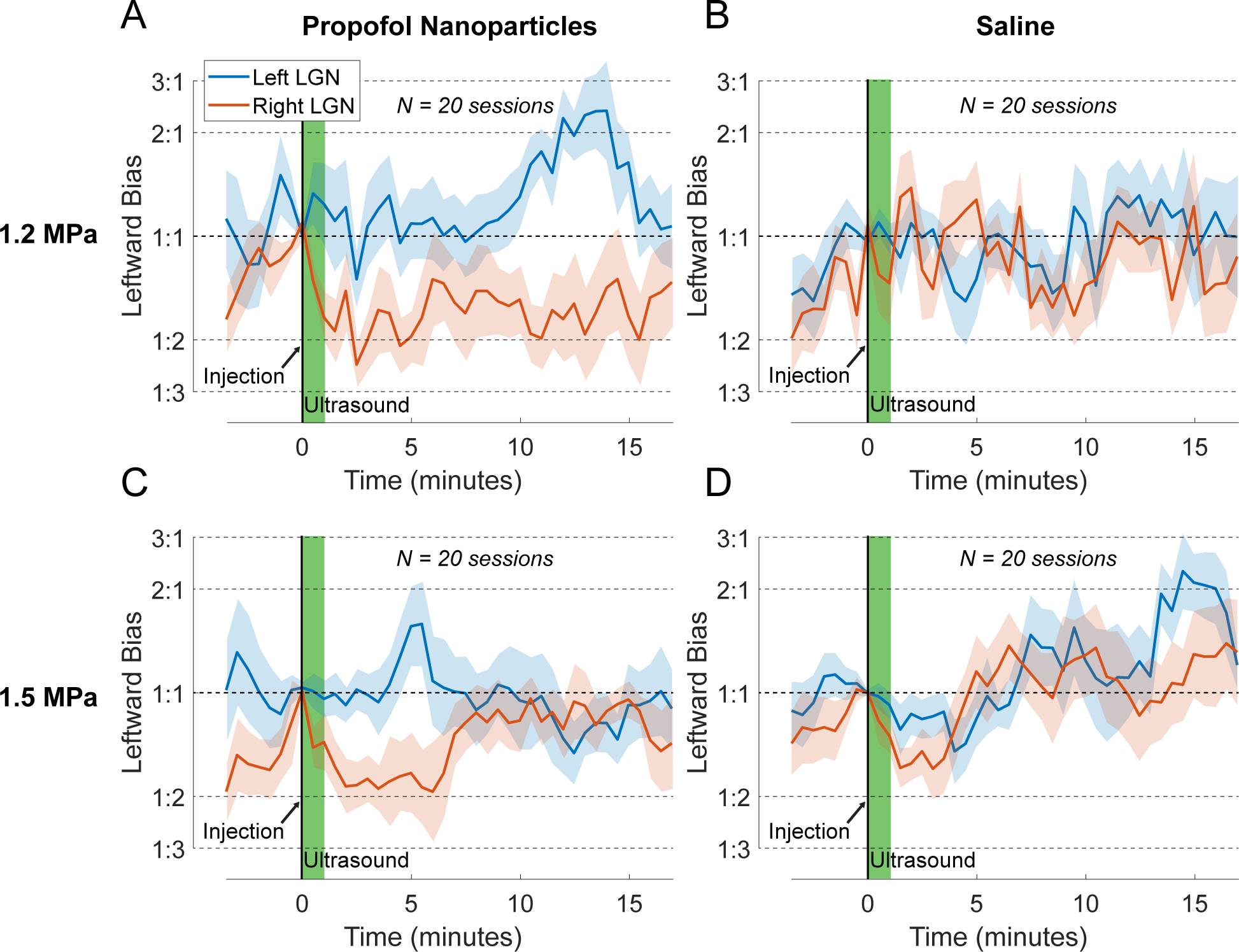
Propofol-induced modulation of choice behavior is target-specific. Same evaluation and format as in Fig. 2, for ultrasound delivered into the right LGN (red) or the left LGN (blue), for the propofol-loaded nanoparticles (A, C) or saline (B, D), and for applied ultrasound pressure of 1.2 MPa (A, B) or 1.5 MPa (C, D). Each plot comprises *n* = 10 sessions per LGN.

### Integrity of the blood-brain barrier

With respect to brain applications, the nanoparticle-based approach is designed to deliver drugs that naturally pass the blood-brain barrier (BBB) without perturbing it. We evaluated the integrity of the BBB following the release using an established approach—gadolinium-enhanced MRI imaging. Specifically, T1-weighted gadolinium-enhanced images have been used to detect the disruption of the blood-brain barrier [35–38]. A 20% signal change is highlighted in Fig. 4A and provides a threshold below which the blood-brain barrier can be considered intact. Across both animals and two independent sessions, in no case did the signal at the target LGN exceed that threshold (representative slices for both monkeys: Fig. 4A; all slices for both sessions and monkeys: Fig. S8). As a positive control, blood vessels are clearly identifiable (Fig. 4A; arrows). Shortening of T1 relaxation times in T1 maps associated with the presence of contrast were also not detected Fig. S7). Moreover, T2-weighted contrast-enhanced imaging, which is commonly used to assess potential edema [39, 40] also did not reveal signal changes (Fig. 4B, Fig. S9). Together, these analyses suggest that the BBB remained intact and that the one-minute pulsed ultrasound exposure did not induce detectable harm to the target tissue.

**Figure 4.**
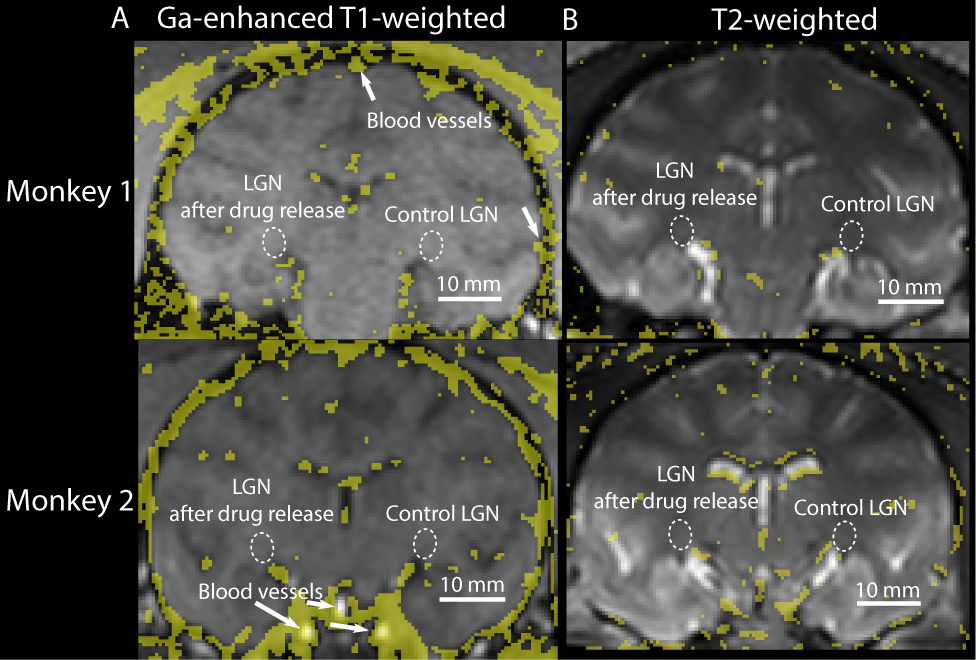
Intact blood-brain barrier following ultrasound-based drug release. Gadolinium-enhanced T1-weighted (left) and T2-weighed (right) MRI contrast images following the administration of propofol-filled nanoparticles (0.5 mg/kg) and one-minute pulsed ultrasound (1.5 MPa) delivered into the left LGN into the two monkeys used in the behavioral task (rows). The ultrasound was delivered using the same hardware and targeting as during the behavioral experiments. Immediately following the release, we injected the contrast agent gadoteridol. The images show the difference before and after the administration of nanoparticles, ultrasound, and gadoteridol. The yellow regions label differences greater than 20% (T1-weighted) and greater than 10% (T2-weighted).

### Pharmacokinetics and biodegradability

We investigated the pharmacokinetics of the nanoparticles in the NHPs. To do so, we incorporated in the nanoparticles a fluorescent dye, along with propofol (see Materials and Methods). We drew blood samples at 2, 10, 20, 40, 80, and 120 minutes following the injection. We then quantified the amount of fluorescence from these blood samples relative to the initial time point. The resulting blood clearance curve (Fig. 5A) shows an initial half-life of 3.1 minutes followed by a slow decay with half-life 195 minutes. This clearance characteristic agrees with those reported using perfluoropentane-based nanoparticles in rats [7, 41]. The dual-exponential nature suggests that the clearance of the nanoparticles involved two distinct processes or organs.

**Figure 5.**
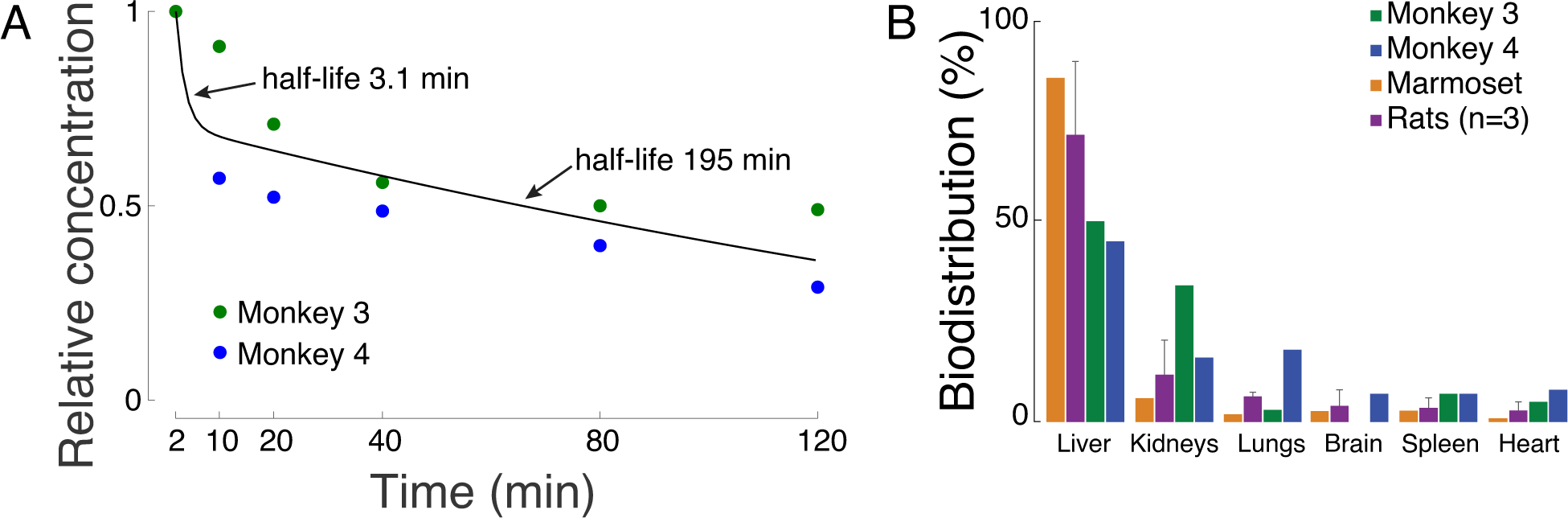
Blood clearance kinetics and organ biodegredation. A) Relative fluorescence as a function of specific sampling times indicated on the abscissa. The nanoparticles were loaded with an infrared dye administered at time 0. The data were fitted with a double exponential function. One exponential showed a fast and the other a slow time constant (see inset). B) Distribution of the nanoparticles in major organs. The figure shows the relative dye accumulation in the respective organs as a percentage of total fluorescence. The rat data are presented as means*±*standard deviation. The brain for Monkey 3 was not available for this analysis.

This pharmacokinetics study relies on the nanoparticles’ ability to retain the infrared dye used for tracking. As shown in Fig. S3, we did not observe any significant leakage of the dye from the nanoparticles. The fluorescence measured from nanoparticles isolated from plasma via centrifugation was consistent over incubation times ranging from 2 minutes to 2 hours. The effect of time in a one-way ANOVA was not significant (*F* (4, 16) = 1.35*, p* = 0.29). Indeed, a previous study [7] has shown minimal serum concentrations of this dye after encapsulation, further suggesting that the leak rate is much slower than the rate of nanoparticle clearance.

We thus evaluated in which organs the PFOB-based nanoparticles degrade. Two macaques, one marmoset, and 3 rats were sacrificed two hours after the injection of dye-loaded nanopartices, and their major organs were extracted for analysis. The majority of the dye-loaded nanocarriers were found in the liver, again in line with previous studies in rats [7, 41]. Appreciable amounts were also detected in the kidneys and lungs (Fig. 5B). These results would be unlikely if significant quantities of dye had leaked from the nanoparticles, as a previous study has demonstrated that free IR800RS dye does not substantially accumulate in the liver [42].

### Clinical chemistry and hematology

The vascular access ports enabled us to repeatedly draw blood following the administration of the nanoparticles into the blood stream, and thus conduct detailed clinical chemistry and hematology evaluations. Since the nanoparticles were found to degrade primarily in the liver (Fig. 5B), we evaluated key markers of liver function—alkaline phosphatase (ALP), alanine transaminase (ALT), and aspartate aminotransferase (AST) (Fig. 6A). ALP and AST remained within normal ranges (green). ALT was mildly elevated during the course of the study and returned to baseline levels in both animals after a two-week period.

**Figure 6.**
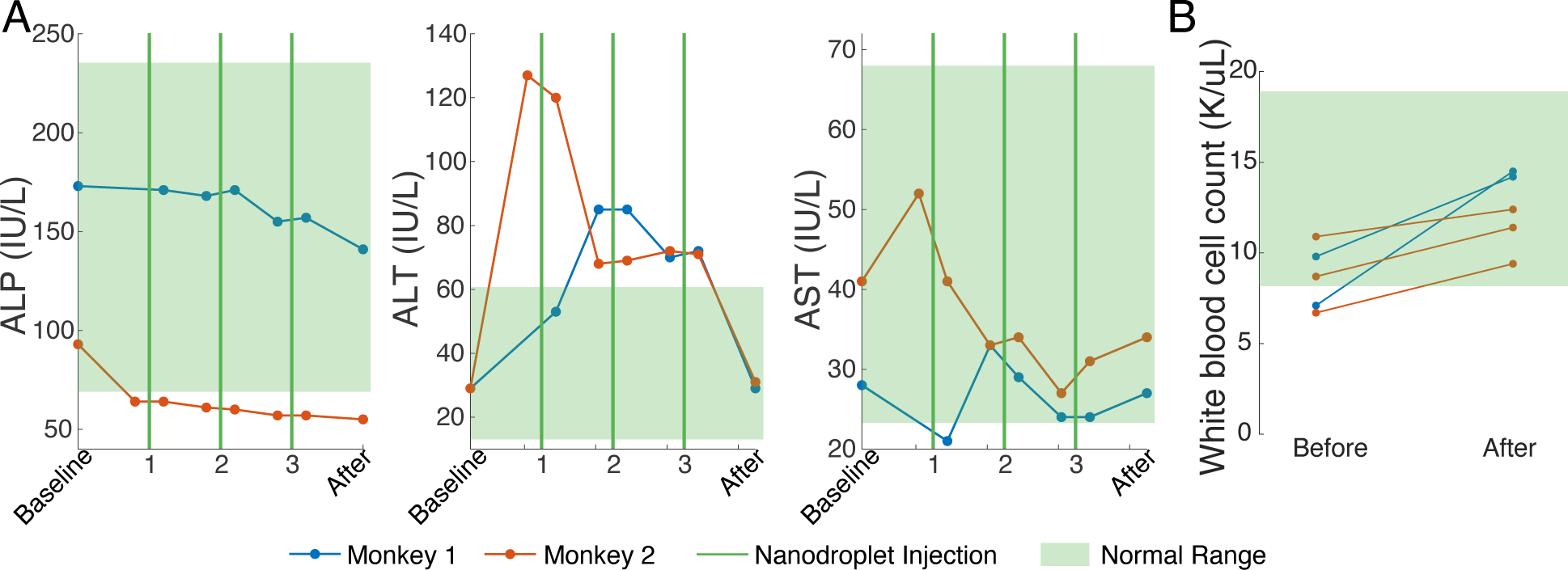
Clinical chemistry and hematology. A) Liver function-relevant serum chemistry: alkaline phosphatase (ALP), alanine transaminase (ALT), and aspartate aminotransferase (AST), across 3 independent sessions. Green lines indicate the time points of the nanoparticle administration. Blood draws were completed immediately before and 1.5 hours after for each administration. For Monkey 1, a preinjection draw in the first session was not available. The first injections for each animal used nanoparticles with propofol concentration of 1.0 mg/kg dose, followed by 0.5 mg/kg for subsequent injections. Baseline was taken prior to any intervention and the final measurement was taken two weeks after the last nanoparticle dose. B) White blood cell counts before and 1.5 hours after the administration.

To assess the response of the immune system, we also evaluated the white blood cell count (Fig. 6B). A detected increase in the white blood cell count was well within the normal range (green). Complete clinical chemistry and hematology analyses are provided in Tables S1 and S2.

## Discussion

This article finds that stable nanoparticles with a high boiling point core can be used to safely release propofol in specific deep brain regions of NHPs. The release of propofol was substantial in that it manifested in specific changes in visual choice behavior. The effect pointed in the expected, neuroinhibitory direction and was reversible. The release is found to be safe at the behavioral, anatomical, and hematological levels. At the behavioral level, we found that the release in visual regions did not impair NHPs’ ability to successfully perform visual discrimination. At the anatomical level, there was no detectable disruption of the NHP blood brain barrier. At the hematological level, a key marker of immune reaction—white blood cell count—remained within normal values. The average half life of the nanoparticles in the NHP blood was found to be about 30 minutes, which provides a practical time window for applications in humans.

A hallmark of this remotely-controlled approach is its ability to flexibly deliver a drug into spatially circumscribed regions of interest. Our finding of target-specific effects on choice behavior (Fig. 2) supports the notion of a spatially-specific release. Specifically, the centers of the left and right LGNs are separated by about 22 mm [43]. Therefore, the approach used in this study has a spatial precision of at least that order. Previous studies in rodents have showed that achieving a sub-cm precision is feasible [8, 11]. In addition, the effect was specific to propofol-filled nanoparticles; saline administered with ultrasound of the same parameters produced no significant effect.

Another key feature of this approach is the low systemic dose of drug required to elicit strong local effects. Anesthesia of macaques typically requires around 2 mg/kg of propofol when administered as a bolus [44]. In comparison, the dose used in our study was just 0.5 mg/kg. This indicates that the approach can be used to release a drug at a target at a high concentration while maintaining a relatively low systemic level. The approach may thus improve current systemically toxic or harmful treatment regimens, such as chemotherapy or treatments of brain circuits with psychedelics.

For brain applications, the nanoparticle-based approach for localized drug delivery differs fundamentally from a related approach, which uses ultrasound combined with blood-circulating microbubbles to transiently disrupt the blood brain barrier (BBB) [45–47]. The nanoparticle-based approach aims to deliver drugs that naturally pass the BBB while preserving its integrity (Fig. 4). Moreover, in the nanoparticle-based approach, drugs are encapsulated within the nanoparticles and shielded by a polymeric shell (Fig. 1B). This design prevents the drug from interacting with tissues and organs until exposed to ultrasound or broken down. In comparison, in the microbubble-based approach, a full dose of a systemically injected drug immediately interacts with all vascularized tissues, thus limiting the specificity of that approach. It is possible that, in combination with microbubbles, these nanoparticles could be used to deliver drugs which do not on their own cross the blood brain barrier. This could be beneficial for chemotherapeutic drugs, which are frequently unable to reach tumors in the brain and could benefit from being localized to the tumor. Nonetheless, such applications should be preceded by careful studies of the interaction between the microbubbles and the nanoparticles, to ensure that appropriate ultrasound parameters are selected to activate the nanoparticles while not causing inertial cavitation of the microbubbles.

All components of the nanoparticles used in this study have been used in humans, which is expected to facilitate regulatory approval. PFOB is well tolerated by humans [48–50], and due to its oxygen-binding capacity has been used as a blood substitute [22–24]. Moreover, the hydrophobic block of the copolymer used—the polylactic acid (PLA)—is generally recognized as safe by the FDA and has been used broadly in modern medicine [51, 52]. Likewise, polyethylene glycol (PEG) has been used extensively for its ability to circumvent activation of the immune system and thus extend the lifetime of drug carriers in the circulation [53, 54]. Indeed, we found that our PEGylated nanoparticles were present in the blood two hours following administration (Fig. 5A). The predominant breakdown of the nanoparticles in the liver suggests an engagement of the reticuloendothelial system [55]. The detected elevations in alanine aminotransferase were below the levels that could indicate hepatocellular damage [56]. Furthermore, the initial increase dropped by the second and third injection and returned to normal following a two-week washout [57, 58].

The approach reported in this study builds on ultrasound-responsive nanoparticles initially developed for localized chemotherapy in rodents using perfluoropentane (PFP) [12, 26] and perfluoro-15-crown-5 ether (PFCE) drug carriers [5]. PFP has a boiling point below the body temperature and thus has raised safety concerns. PFCE has a much higher boiling point—and thereby stability—but its biological half-life of nearly 8 months limits clinical deployment [59]. The perfluorocarbon core used in this study, PFOB, addresses both issues, but difficulties have remained in regard to its relatively high activation pressures [60], as activation pressure appears to correlate with the boiling point [61, 62]. We have resolved this issue using relatively low ultrasound frequencies (248-480 kHz). In this ultrasound frequency range, we observed effective release *in vitro* (Fig. S1A) and *in vivo*, finding a strong and selective modulation of choice behavior of NHPs (Fig. 2, Fig. 3).

While low boiling point PFC nanoparticles have typically been activated by vaporization of the core itself, this is unlikely to occur for PFOB given its 142*^◦^*C boiling point. Thus, drug release in this case is likely mediated by a mechanical effect, which may include particle displacement or acoustic radiation force. Indeed, similar perfluorocarbon-based nanoparticles have been shown to effectively release drug *in vivo* without evidence of inertial or stable cavitation [26]. It is possible that the mechanical effects are driven or enhanced by the mismatch between the acoustic impedance of the blood (1.65 MRayl) and the perfluorocarbon nanoparticle core (1.84 MRayl [63, 64]). Mismatched acoustic impedance can enhance drug release without resting on cavitation [65]. Nonetheless, more work is needed to elucidate the mechanisms of drug release.

This study has two limitations. First, although the study uses an established behavioral framework and the well-characterized visual system to evaluate neuromodulatory effects [27–33], there is a lack of a modality to image the released propofol. Microdialysis was not used due to concerns of a needle disrupting the blood-brain barrier, which would artificially boost the release. Second, the reported behavioral effects are subject to adaptation and potentially other higher-order cognitive influences [28]. This issue is mitigated by contrasting the release across the two brain sites and by contrasting propofol-filled nanoparticles with saline.

It is possible that the volume of the drug release may exceed the ultrasound focus. On this front, propofol and other hydrophobic small molecule drugs have a strong propensity to diffuse into the brain. Propofol diffuses rapidly out of the bloodstream and into the brain, with a blood-brain equilibration half-life of just 2.9 minutes when administered systemically [66]. This may explain why a previous study found a highly confined release of propofol by ultrasound in the brain [8].

These NHP data are expected to lead to an accelerated adoption of this targeted approach in humans. For instance, the propofol-containing nanoparticles could be used for virtual lesioning of individual candidate regions involved in epilepsy, pain, or mental disorders. Longer-acting drugs could subsequently be delivered into the identified malfuncioning circuits and thus provide personalized, targeted therapy. Targeted delivery of drugs for other indications such as cancer, and organs other than the brain, could fuel additional important applications.

## Conclusions

In summary, we have developed an approach that enables ultrasound-triggered delivery of drugs into circumscribed regions in the brain or the body as demonstrated by robust modulation of choice behavior and a favorable safety profile in NHPs. This targeted pharmacomodulation approach has the potential to provide treatments to individuals for whom current drug treatments cause unbearable or undesirable side effects.

## Materials and Methods

### Materials

Methoxy poly(ethylene glycol)-b-poly(D,L-lactide) (PEG-PDLLA) co-polymers with 2: 2.2 kDa molecular weights, respectively, were obtained from PolyScitech (USA). Perfluorooctyl bromide were obtained from Tokyo Chemical Industry Co. (Japan). Propofol was obtained from Sigma Aldrich (Millipore Sigma, Canada). Infrared dye IR800RS NHS Ester was obtained from LI-COR Biosciences (USA). HPLC-grade tetrahydrofuran (THF), n-hexane, and methanol were obtained from Fisher Scientific (USA). Sterile phosphate buffer solution (PBS) was obtained from Cytiva Life Sciences (USA). Pooled, blood derived human plasma with anticoagulant K2 EDTA was obtained from Innovative Research (USA).

### Animals

All procedures conformed to approved protocols by the Animal Care and Use Committee of the University of Utah (protocols 18-11011, 18-12015, and 21-12012). For all behavior studies and safety studies not requiring euthanasia, we used two male rhesus macaques (*macaca mulatta*) with weights 10.0 kg and 13.8 kg, both aged 8 years (monkeys 1 and 2). Two adult male rhesus macaques (*macaca mulatta*, monkeys 3 and 4), an adult male common marmoset (*callithrix jacchus*), and 3 adult male Sprague-Dawley rats participated in the safety and pharmacokinetics experiments. These macaques weighed 10.8 and 16.1 kg (age 6 and 13 years). The marmoset weighed 288 g (age 7 years) and was from an in-house colony. The rats weighed 830-900 g (age 1 year) and were obtained from Charles River. Rats were selected for the initial safety screening because they are more easily available in larger quantities. The marmoset and the first two macaques became accessible upon successfully completing their intended studies in our laboratory and those of our collaborators. This minimized the risk to animals still participating in ongoing studies.

The macaques were preferentially housed in pairs, according to the approved protocol. They were given daily enrichment by husbandry staff and received fruit and vegetables daily to supplement their diet.

The behavioral experiments were performed in monkeys and were not terminal. Rats were selected for the initial safety screening as they are more suitable for terminal experiments. The marmoset and the first two macaques became available for the terminal experiments following the completion of their original studies. All procedures were IACUC-approved.

### Nanoparticle production

The process of manufacturing the drug-encapsulating, ultrasound-responsive PFC particles is described in detail in previous studies [12, 26]. The process converts small (tens of nanometers in diameter) micelles into much larger (hundreds of nanometers) PFC nanoparticles. First, the PEG-PDLLA polymer constituting the basis of the nanoparticle shell was dissolved in THF at a ratio of 1mL THF: 20 mg polymer. For the biodistribution and blood clearance studies, infrared dye was added at a ratio of 1:32 (dye:polymer) for the rats and marmoset and 1:110 or 1:89 for monkeys 3 and 4, respectively. The macaques used disproportionally smaller quantities of the dye than the smaller animals for supply reasons. THF was then allowed to evaporate overnight until a gel-like layer remains. PBS was added at a ratio of 1 mL PBS: 20 mg polymer and placed on a shaker table at 120 rpm to dissolve for 15 minutes. The addition of PBS orients the hydrophilic copolymer, PEG, toward the water and the hydrophobic copolymer, PDLLA, away from the water, forming micelles. Next, the PFOB core and drug were added and emulsified. A 2:1 ratio of polymer to drug was used. The nanoparticles’ diameter can be controlled by the ratio of PFC to polymer, as reported previously [5]. We used a ratio of 3 *µ*L PFOB: 1 mg polymer for all experiments except the 1.5 MPa behavior studies, for which the ratio was adjusted to 2.75: 1. The PFC and drug were added to 15 mL centrifuge tubes and gently shaken to combine before adding 8 mL of the micelle solution. A 20 kHz, 500W sonicator with a cup horn attachment (VCX500, Sonics & Materials, Inc., USA) emulsified the PFOB and drug, forming stable nanoparticles. The samples were sonicated in a 10*^◦^*C water bath at 20% power for 3 minutes, inverting the tubes halfway through to ensure even distribution. A custom temperature-controlled cooling system maintained the bath temperature during sonication. We found this controlled temperature approach to maximize the consistency of the nanoparticle sizes, drug encapsulation, and release properties. The resulting solution contained the desired nanoparticles in addition to remaining micelles, dissolved polymer, and free propofol. Nanoparticles were isolated from micelles and free drug using three cycles of centrifugation at 3,000 g relative centrifugal force (RCF) at 4*^◦^*C. After each cycle, the supernatant was discarded and the pellet dissolved in fresh PBS. Blank nanoparticles were manufactured using the same process but no propofol is incorporated before sonication. The concentration was estimated based on the average encapsulation rate of propofol-loaded particles.

### Nanoparticle characterization

Nanoparticle sizes were measured using a Zetasizer Nano S (Malvern Panalytical, UK), which reports the intensity-weighted size distribution. The size values reported in the Supplementary Materials section describe the z-average diameter *±* standard deviation of the distribution of the intensity values measured by the device. To quantify the amount of drug encapsulated, 25 *µ*L of nanoparticle solution was added to 225 *µ*L of methanol to dissolve all components. A UV-vis spectrophotometer (NanoDrop 2000, Thermo Scientific, USA) was used to quantify the concentration by comparing the absorbance at 276 nm to a propofol standard curve.

Drug release efficacy was assessed *in vitro* using a solvent extraction method similar to that described in [7]. In this procedure, 0.2 mL nanoparticle solutions in either human plasma or PBS are placed in contact with 0.2 mL of an organic sink (hexane) in a 1.5 mL centrifuge tube while ultrasound is applied from a single-element transducer coupled to the tube with degassed water. Following sonication, the hexane was extracted to measure the concentration of propofol released using UV-vis spectrophotometry. The time hexane was in contact with the nanoparticles was held constant at 105 seconds to control for the time course of drug release. Percent drug release is reported as the amount of drug released into hexane relative to the drug encapsulated. The mean*±*SD percentage of drug encapsulated across all behavior studies was 10.6*±*1.9 %.

To assess the stability of nanoparticles in human plasma, the nanoparticles were first dissolved in the plasma following the last centrifuge cycle. Drug release with and without ultrasound was quantified as described above immediately, after 1 hour, and after 2 hours of incubation with plasma. Ultrasound pressure was held constant at 1.5 MPa.

To quantify the rate of dye leak *in vitro*, nanoparticles were prepared using 0.5 mg of IR800RS dye, 20 mg of propofol, 55 *µ*L of PFOB, and a micelle solution containing 40 mg of PEG-PDLLA polymer. After the final centrifuge cycle, the nanoparticle pellet was dissolved in plasma and incubated for either 2, 20, 40, 80, or 120 minutes. Then, the solution is centrifuged again for 5 minutes and 3,000 g RCF to isolate the nanoparticles in the pellet. The pellet was then dissolved in 0.1 mL of PBS and fluorescence of the dye quantified using IVIS imaging (see IVIS imaging subsection for details).

### Sterilization and endotoxin testing

For animal studies, nanoparticles were sterilized with either ultraviolet light or filtration. UV light was selected for the acute studies for its efficacy of sterilization while it is considered unlikely to disrupt polymeric drug carrier structure and function [67]. Samples in glass vials were exposed to an 8W UV lamp (Philips, USA) in a custom chamber for 3 hours. For the longer-term studies in monkeys, we sterilized the micelle solution by 0.2 micron filtration, then used sterile reagents for the remainder of the production. Filtration of the finished product was not possible because the nanoparticles are mostly larger than the filter pore size.

To confirm that the process does not introduce unwanted contamination, we tested for the presence of bacterial endotoxin, an FDA-required step for any drug product [68]. We used gel-clot lyophilized amebocyte lysate tests (ToxinSensor, Genscript, USA) to detect the presence of endotoxin above the detection threshold. This semi-quantitative method was used to ensure that the nanoparticles contained a level of endotoxin lower than the FDA allowable limit of 5.0 EU/kg when diluted at doses to be administered to animals.

### Nanoparticle dosing

Long-term study animals were implanted with vascular access ports to enable rapid intravenous infusions and blood draws. The vascular access port system consists of the port itself and a polyurethane catheter (Swirl-Phantom and Hydrocoat, Access Technologies, USA) inserted via the saphenous vein into the inferior vena cava. Dosing was ramped up over four sessions to minimize potential safety issues: 0.1, 0.33, 0.66, and 1.0 mg/kg of propofol. We chose to use a dose of 0.5 mg/kg, at which animals showed no signs of drowsiness.

### Behavior experiments

The system for delivering ultrasound to the lateral geniculate nucleus and monitoring behavior effects is described in previous publications [27, 28] and illustrated in Fig. 1. Two non-human primates were trained in a visual discrimination task which has commonly been used in neurology and neuroscience [27, 28, 30, 69]. In this task, the subject fixates on a central target presented on a screen as detected using an eye tracker (Eyelink, SR Research Ltd., Canada). After a randomized delay, the fixation point disappears and a target is presented in either the left or right visual hemifield. Following another delay, a second target is presented in the opposite visual hemifield. The subject is rewarded with a probability between 50% and 100% with a drop of juice from a lick spout for selecting the target presented first. In each session, there were 5 possible delays between the onset times of the two targets, with the delays calibrated to each subject. Delays are selected such that the subjects are near 100% accuracy in the longest delays and near 50% accurate at the middle delay (0 ms), with some degree of uncertainty at the delay in between. Using these delays, we fit a sigmoid function to each subject’s behavior to quantify the magnitude and the polarity of neuromodulatory effects.

In this study, we quantified behavior by fitting a sigmoid curve to 3 minute segments of data. The subjects first performed at least 3 minutes of behavior before any intervention to establish a baseline. We use this baseline behavior to establish the point of equal preference, which is the time delay at which the sigmoid curve crosses the 50:50 choice proportion line. Animals were not cued to perform the task while we administered propofol-loaded nanoparticles or saline. The task performance was resumed upon the delivery of the ultrasound. In 3-minute segments following sonication onset, we measured effects on choice behavior as the percent of leftward choices at the delay of equal probability established during the baseline. A timeline of the experiment is shown in Fig. 7. Plots in Fig. 2 are an average of results from 5 sessions for each condition for each subject, for a total of *n* = 80 sessions. One saline session was excluded due to a lack of trials in the 3 minute baseline period. We interleaved sessions with nanoparticles and saline and the sonicated side. Sessions were completed on a nearly daily basis.

**Figure 7.**
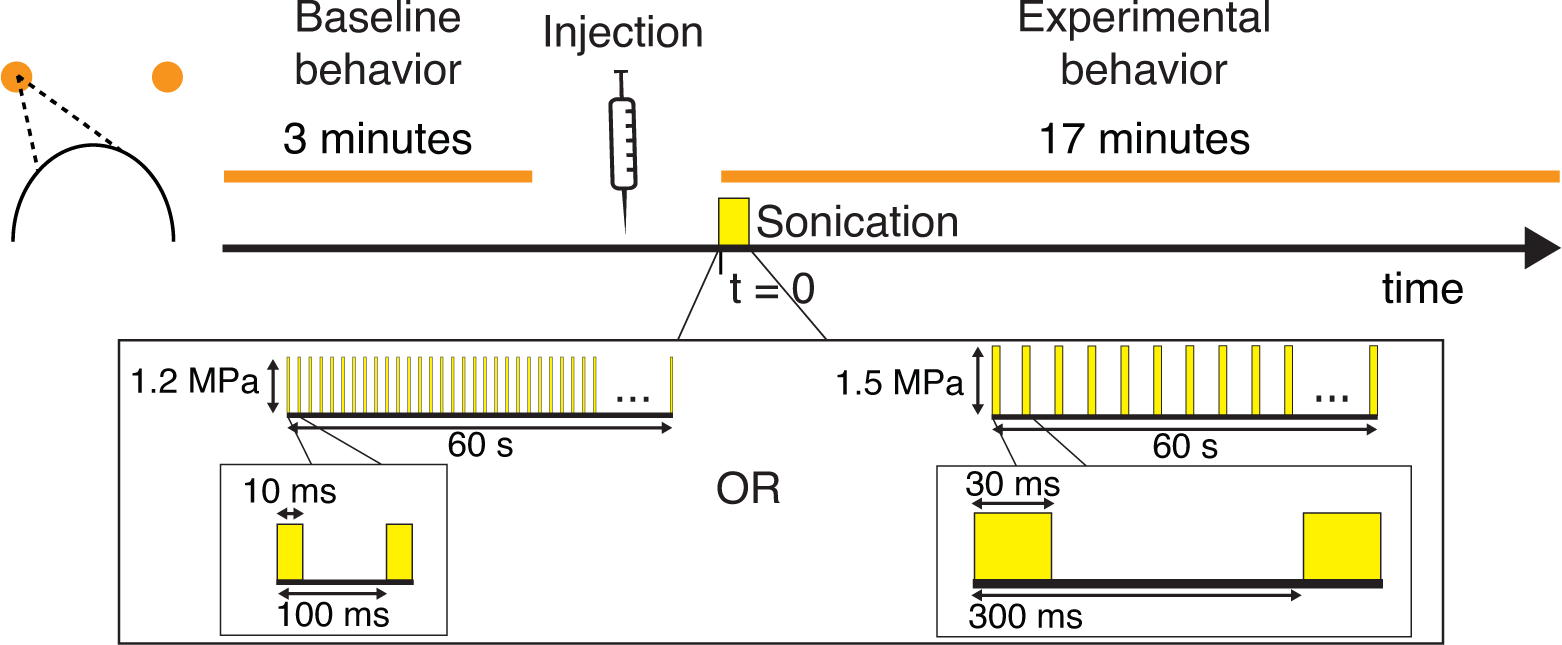
Behavior Experiment Timeline. Each behavior session began with the animal performing the task for at least 3 minutes to establish a baseline. The task was paused while the experimenter administered nanoparticles or saline, then resumed at the time of ultrasound onset. The animals completed at least 17 minutes of the task after ultrasound onset.

### Statistical Analysis

Statistics are calculated in the behavior study over number of repeated sessions rather than number of animals. Statistics were assessed over the individual sessions. The primary analysis is four-way ANOVA with factors of LGN sonicated (left/right), pressure (1.2/1.5 MPa), type of injection (propofol nanoparticles/saline), and subject (Monkey 1/2). Five sessions were completed by each animal for each condition, resulting in 80 sessions in total (left and right sonication, propofol nanoparticle and saline injection, 1.2 and 1.5 MPa sonication). The time window from 2-5 minutes following the ultrasound onset was selected for the analysis as it was the peak in behavior bias observed in the time courses. The ANOVA was computed in MATLAB for all the behavior sessions with saline and propofol (Fig. 2), then repeated after splitting the data into 1.5 MPa and 1.2 MPa sonications (Fig. 3). We also have quantified these effects for each animal separately using the same ANOVA design but without the factor of subject. In order to further confirm the statistical robustness of these effects, we repeated this procedure with blank nanoparticles at 1.5 MPa - no propofol was incorporated (Fig. S4). The two-way ANOVA was then repeated between each of these conditions and a Bonferroni correction applied for the three comparisons. Box plots of the data analyzed are available in Fig. S5. A one-way ANOVA was used to quantify the effect of ultrasound pressure on drug release *in vitro* from 0 to 2.5 MPa.

### Pharmacokinetics studies

The rats were anesthetized with 2.5-3% isoflurane, and dye-loaded nanoparticles were administered at a dose of 1 mg/kg propofol followed by an equal volume of sterile saline. After one hour, the animals were euthanized by exsanguination under 5% isoflurane anesthesia.

The primates were preanesthetized with ketamine (25 mg/kg intramuscularly) and intubated with endotracheal tubes. They were artificially ventilated and anesthesia maintained with 1-4% isoflurane throughout the procedure by veterinary staff. The animals were placed on heated operating table pads to maintain body temperature. For the marmoset, dye-loaded nanoparticles were injected through the tail vein at doses of 1 mg/kg propofol for each of two injections separated by 45 minutes. A total volume of 2 ml of nanoparticle solution was administered, followed by an equal volume of sterile saline. The marmoset was euthanized by an overdose of sodium pentobarbital and perfused transcardially with 4% paraformaldehyde 82 minutes after the first injection. For Monkey 3, one injection of dye-loaded nanoparticles was administered in the right saphenous vein at 1 mg/kg propofol and a volume of 5 mL, followed by an equal volume of sterile saline. Blood samples were taken from the left saphenous vein 2, 10, 20, 40, 80, and 120 minutes, at a volume of 1 mL each. For Monkey 4, the right (injection) and left (blood draws) cephalic veins were used with a 1 mg/kg propofol dose and 10 ml volume of dye-loaded nanoparticles. Following 120 minutes of monitoring, the monkeys were euthanized by an overdose of sodium pentobarbital and perfused transcardially with 4% paraformaldehyde.

The procedures were painless and wholly consistent with the recommendations of the Panel on Euthanasia of the American Veterinary Medical Association.

### IVIS imaging

Major organs were extracted and subjected to infrared fluorescence analysis using an IVIS system (Perkin Elmer, USA) with methods similar to those previously demonstrated [7]. Excitation was set to 745 nm and emission to 820 nm, with 5 seconds of exposure. 2 seconds of exposure was used for the *in vitro* samples. Extracted organs were placed directly on a dish in the field of view of the camera. For monkeys, the same organs as in the smaller animals were available, with the exception of the brain for Monkey 1. Blood samples were pipetted into drops of 100 uL on a dish. Total fluorescence for each of these samples was quantified using Aura software (Spectral Instruments, Inc., USA), defining regions of interest incorporating the whole organ or blood sample and subtracting a background region of the same size. Percent biodistribution was computed as the total fluorescence of the region containing the organ divided by the sum of the fluorescence of all organs. Nanoparticle concentration in the blood was computed as the amount of fluorescence from each sample relative to the first sample, obtained at 2 minutes.

### Ultrasound parameters

Ultrasound parameters for all experiments are summarized in Table 1. The ultrasound carrier frequency in the *in vitro* drug release experiments was 300 kHz using a focused single element transducer (H-115, 64 mm diameter, 52 mm focal depth, Sonic Concepts, USA). Stimuli were generated using a function generator (33520b, Keysight Technologies, USA) and the signal amplified using a 55-dB, 300 kHz–30 MHz power amplifier (A150, Electronics & Innovation, Ltd., USA). Pulses 100 ms in duration were repeated once per second for a total of 60 seconds. This pulsing sequence was previously used to activate PFC nanoparticles in rodents [7, 26]. The pressure levels at the vial location, measured in degassed water, were 0, 0.5, 0.9, 1.3, 1.7, 2.1, and 2.5 MPa. The pressure fields were measured using a capsule hydrophone (HGL-0200, Onda Corporation, USA) calibrated between 250 kHz and 40 MHz and secured to 3-degree-of-freedom programmable translation system (Aims III, Onda). The single-element transducer has a half-power beamwidth of 6 mm x 40 mm.

**Table 1.** Ultrasound parameters for all experiments. The 300 and 248 kHz sonications were delivered by the single-element H-115 transducer, while the 480 kHz sonications were delivered by the 256-element array.

| Experiment | Animal(s) | US Frequency | Pressure | Pulse Length | Pulse Frequency | Duration |
| --- | --- | --- | --- | --- | --- | --- |
| Behavior | Macaques 1 and 2 | 480 kHz | 1.2 MPa | 10 ms | 10 Hz | 1 min |
| Behavior | Macaques 1 and 2 | 480 kHz | 1.5 MPa | 30 ms | 3.33 Hz | 1 min |
| MRI | Macaques 1 and 2 | 480 kHz | 1.5 MPa | 30 ms | 3.33 Hz | 1 min |
| Pharmacokinetics | Macaques 3 and 4, Marmoset, Rats 1-3 | 248 kHz | 1 MPa | 100 ms | 1 Hz | 1 min |
| *In Vitro* | n/a | 300 kHz | 0-2.5 MPa | 100 ms | 1 Hz | 1 min |

In the pharmacokinetics experiments, the same transducer was operated at 248 kHz in 60-second blocks of 100 ms pulses. The minimum operating frequency was selected as a compromise between high release effectiveness and small release volume. The animals were shaved and the transducer coupled to the scalp using a 2% agar cone and ultrasound gel. The maximum pressure was estimated to be 1 MPa for all animals. The pressure was estimated from free-field measurements and a correction for the skull which indicates a transmission rate of 76% through the skull of a 780g rat [70] and 66% through a macaque skull [71]. Transmission through the marmoset skull has been less thoroughly studied but was predicted to be similar to rats since the species are comparable in weight. Ultrasound was applied to the rats at the midline, 2 mm posterior to the eyes for 60 seconds 5 minutes after administration of nanoparticles. For the marmoset, ultrasound was applied at the posterior surface of the skull for 60 seconds 5 minutes after each of the two administrations. The right and left visual cortex were targeted independently for Monkey 3. Ultrasound was applied in 60-second blocks over 90 minutes for a total of 4 sonication blocks per side starting 2 minutes after nanoparticle administration. Sonication of the left and right sides was interleaved. For Monkey 4, the right and left visual cortex were targeted simultaneously using two ultrasound transducers to maximize the sonicated volume. In this monkey, sonications were repeated for two 60-second blocks separated by two minutes. No ultrasound was delivered after this period due to initial results which indicated that the nanoparticle concentration in the bloodstream decays rapidly (Fig. 5A).

In the behavioral, MRI, and blood draw components of the investigation, we used a custom 256-element phased array (Guangzhou Doppler Electronic Technologies, China) that allows us to target the deep brain - specifically, the lateral geniculate nuclei, as described in detail in our previous publications [27, 28]. This transducer has a half-power beamwidth 1 mm x 3.75 mm [27, 28]. Briefly, the scalp was shaved before each session and the transducer affixed to the animal’s head via implanted titanium pins (Gray Matter Research, USA) and coupled using a 6% polyvinyl alcohol (Fisher Scientific, USA) cryogel. The transducer was driven by the Vantage256 controller (Verasonics, USA) at 480 kHz. This corresponds to the lower corner frequency of the transducer’s bandwidth and was selected to maximize the mechanical effects on the nanoparticles. Ultrasound pressure and focal location were determined using MR thermometry. Details on this procedure are published [27, 28]. Ultrasound at the frequency of 480 kHz was pulsed at 10% duty cycle for one minute for all experiments. In the behavioral experiments, we delivered into the targets a peak pressure of 1.2 MPa in 10 ms pulses or 1.5 MPa in 30 ms pulses. The 1.5 MPa sessions were completed with a longer pulse duration as previous studies have indicated that this can lower the activation threshold of PFC nanoparticles and enhance drug release [7, 10].

### MR imaging

All studies were performed on a 3T Vida MRI scanner (Siemens, Germany). The MR imaging was used to assess the potential disruption of the blood-brain barrier (BBB) or tissue damage. Two sessions of propofol-based nanoparticle release were performed for Monkey 1 and one for Monkey 2 under the MRI. Potential disruption of the BBB is assessed using gadolinium-enhanced T1-weighted imaging, which is a common approach since gadolinium does not cross an intact BBB [35, 72, 73]. T2-weighted images were taken to inform on a potential edema formation [37]. We followed a previous NHP protocol to evaluate these effects [35]. These methods were established to evaluate the volume of potential blood brain barrier disruption [35, 74, 75]. In each experiment, we first collected a T1-weighted scan (3D Volumetric Interpolated Breath-hold Examination (VIBE), TR/TE=4.46/1.42 s, 192×132×80 mm field of view, 1 mm isotropic voxels, readout bandwidth 490 Hz/pixel, 5 averages, 3:55 minutes) and a T2-weighted scan (Sampling Perfection with Application optimized Contrast using different flip angle Evolution (SPACE), TR/TE=4000/179 ms, 192×136×80 mm field of view, 1 mm isotropic voxels, readout bandwidth 789 Hz/pixel, Turbo factor 165, 1 averages, 3:30 minutes) to use as baselines. T1 maps were reconstructed from data acquired with a Short TI Inversion Recovery (STIR) turbo spin echo pulse sequence for one session with Monkey 1. Imaging parameters included TR/TE=8000/8 ms, 192×132 mm field of view, 1×1 mm voxels, 2 mm slice thickness, echo train length 14, readout bandwidth 1002 Hz/pixel, acquisition time 1:30 minutes for each of 8 TIs: 25, 50, 100, 250, 500, 1000, 2500, and 6000 ms. Quantitative T1 mapping has been used previously to detect subtle BBB lesions which are not otherwise apparent in standard radiological practice as the presence of gadolinium contrast can decrease T1 times [76]. We then administered a solution of nanoparticles at a concentration of 0.5 mg/kg of propofol followed by 10 mL of sterile saline via an intravenous catheter placed in the subjects’ arm. Ultrasound was delivered starting two minutes after completion of the nanoparticle injection. Immediately following the ultrasound delivery, gadolinium-based contrast agent was administered at a dose of 0.15 ml/kg (ProHance Gadoteridol, 279.3 mg/mL, Bracco Diagnostics, USA). We administered the contrast agent after the drug delivery procedure to use the same approach as previously [35, 74, 75, 77]. Repeated T1 and T2 scans and T1 maps were taken 30 minutes following contrast administration, which allows gadolinium in larger blood vessels to diffuse away. The monkeys were not moved after the initial imaging sequences began - nanoparticles and gadoteridol were administered via an IV catheter line accessed from outside the scanner and ultrasound was applied via a fixed transducer placed prior to imaging.

MR images were initially processed in MATLAB by normalizing to the average value of an off-target region of the brain 10 mm anterior to the sonication target. Percent change from baseline was determined by dividing the post-treatment image by the baseline image. A change of greater than 20% from baseline was interpreted to indicate presence of gadolinium and therefore BBB opening. This threshold is high enough to circumvent the noise caused by lingering gadolinium in blood vessels [35]. A threshold of 10% has been reported previously [35, 74], but we increased this threshold to reduce the influence of gadolinium in blood vessels and areas with very low initial intensity. T2-weighted images were processed the same way, using a 10% threshold of intensity increase. We analyzed 5 slices (5 mm) in each direction from the location of the LGN, shown in Fig. S8 and Fig. S9.

## Acknowledgements

We thank Dr. Melanie Graham for performing vascular access port surgeries, Dr. Caroline Garrett for veterinary support, Dr. Frederick Federer for assistance with the pharmacokinetics studies, and Dr. Patrick Tresco for assistance with an initial proof of concept.

## CRediT authorship contribution statement

**Matthew Wilson:** Writing - original draft, Conceptualization, Data curation, Formal analysis, Investigation, Methodology, Project administration, Resources, Software, Validation, Visualization. **Taylor Webb:** Writing - review & editing, Conceptualization, Data curation, Formal analysis, Investigation, Methodology, Software, Validation, Visualization. **Henrik Odéen:** Writing - review & editing, Data curation, Investigation, Methodology. **Jan Kubanek:** Writing - review & editing, Conceptualization, Funding acquisition, Investigation, Methodology, Project administration, Supervision, Visualization.

## Data Availability

Data will be made available on request.

## Supplementary Information

### Nanoparticle characterization

The mean*±*SD diameter of the nanoparticles was 473*±*28 nm for *in vitro* drug release experiments, 550*±*83 nm in MRI studies, 809*±*328 nm for behavior studies at 1.2 MPa, and 526*±*109 nm at 1.5 MPa. Blank nanoparticles averaged 986*±*444 nm in diameter. The size distributions of PFOB nanoparticles were found to be stable over a 24-hour time period when stored in PBS at room temperature. The average size distribution for blank and propofol-loaded nanoparticles is shown in Fig. S1B. The somewhat larger average size of the nanoparticles prepared for the behavioral studies is likely due to the higher quantities of all materials necessary for the injection into a large animal. The resulting higher concentration of particles may encourage aggregation.

**Figure S1.**
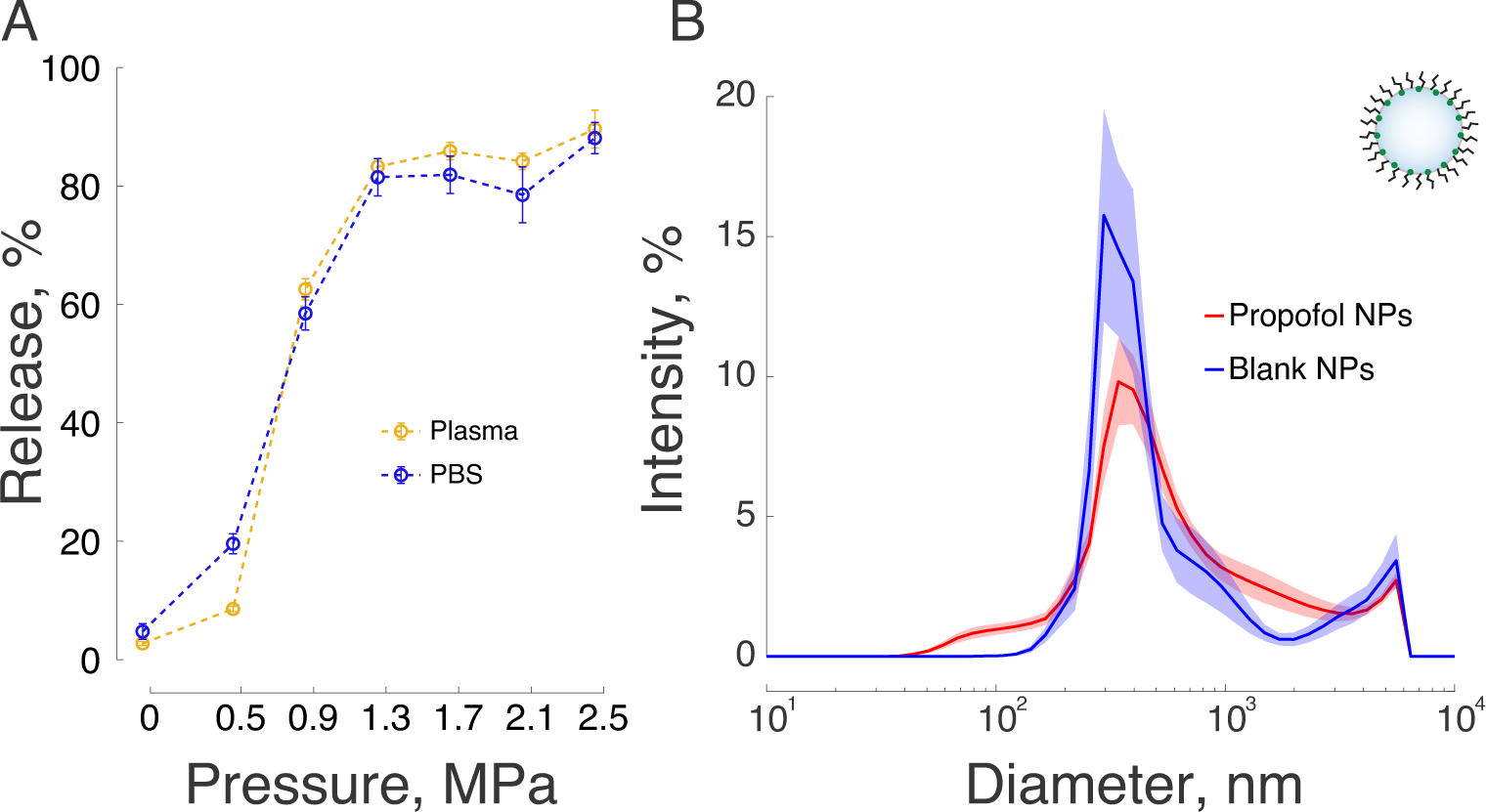
PFOB nanocarriers effectively release propofol in response to ultrasound *in vitro*. A) Mean*±*s.e.m. percentage of propofol released into hexane in response to 300 kHz ultrasound of ultrasound pressure amplitude indicated on the abscissa. Nanoparticles were suspended in either PBS (blue) or human plasma (yellow). The data comprise *n* = 4 independent vials for 0 MPa in PBS and all plasma datapoints, and *n* = 3 for the remaining pressures in PBS. B) Mean*±*s.e.m. size distribution of propofol-loaded and blank nanoparticles for all behavior experiments. *n* = 34 batches of propofol-loaded nanoparticles and *n* = 13 blank.

### Blank nanoparticles

The effects of blank nanoparticles with sonication were indistinguishable from saline injection, indicating that the particles themselves do not contribute to the neuromodulatory effects, but rather the drug that is released. Adding the blank nanoparticles into the analysis of all 1.5 MPa sonication sessions, the interaction of LGN sonicated and drug is still statistically significant (*F* (2, 46) = 7.22*, p* = 0.0019). The difference between blank nanoparticles and propofol was significant (*F* (1, 29) = 8.77*, p* = 0.0061). There was no significant difference between blank nanoparticles and saline (*F* (1, 30) = 0.00*, p* = 0.98). There was a significant interaction of the sonicated LGN side (left or right) and intervention (saline, blank nanoparticles, or propofol-filled nanoparticles) in each monkey (Monkey 1: *F* (2, 21) = 3.75*, p* = 0.041, Monkey 2: *F* (2, 23) = 4.46*, p* = 0.023).

**Figure S2.**
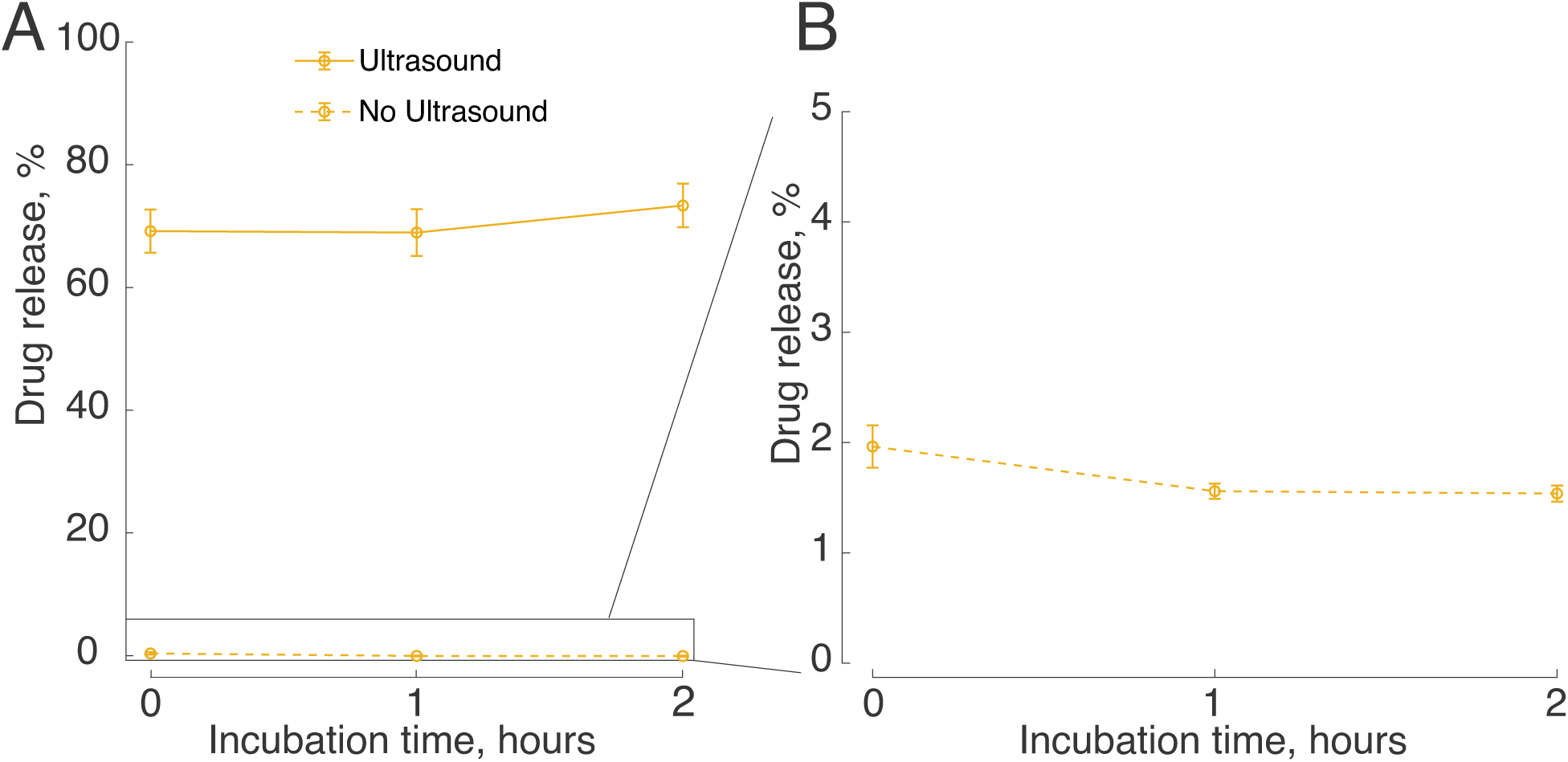
Nanoparticles are stable over two hours in plasma. A) *In vitro* propofol release percentage with 1.5 MPa ultrasound (solid line) and without ultrasound (dotted line). Drug release was not affected after contact with plasma, suggesting the particles remain stable and responsive to ultrasound. B) *In vitro* propofol release percentage without ultrasound.

**Figure S3.**
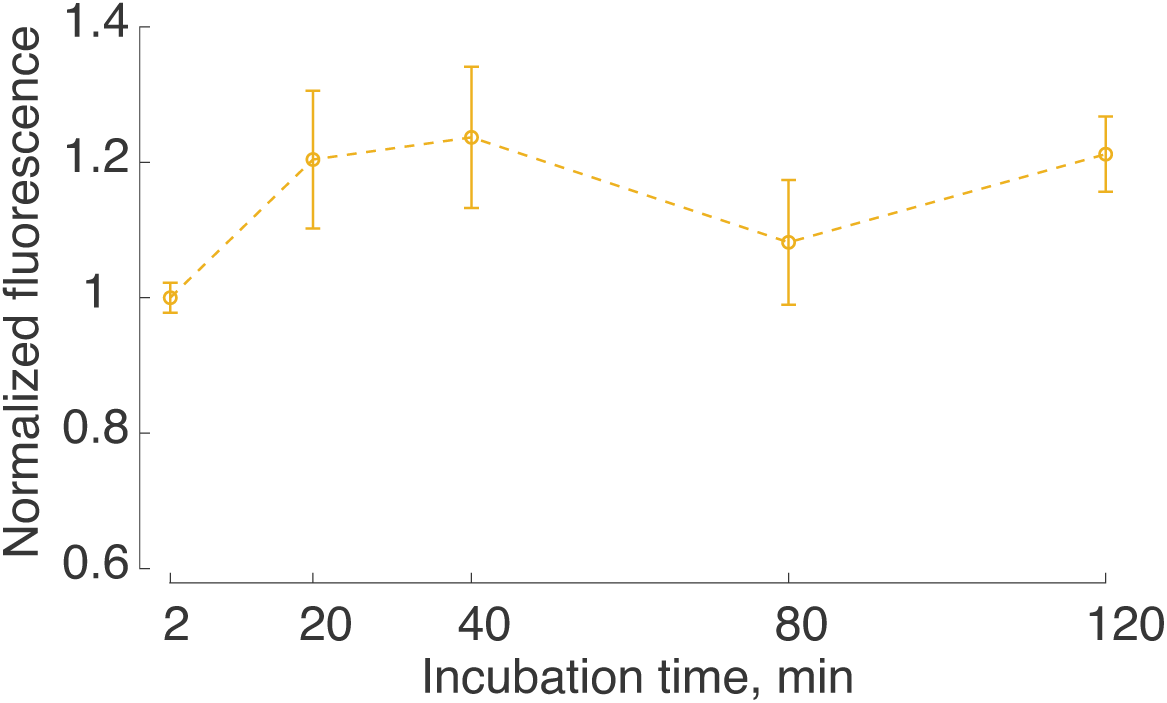
Dye-loaded nanoparticles do not leak dye beyond 2 minutes of contact with human plasma *in vitro*. Dye-loaded nanoparticles were incubated in human plasma for times from 2 to 120 minutes, mirroring the time span of the pharmacokinetics studies shown in Fig. 5. The plasma was then centrifuged to isolate the nanoparticles and the fluorescence of the pellet quantified. Fluorescence for each of *n* = 4 samples per time point were normalized to the mean of the 2 minute samples.

**Figure S4.**
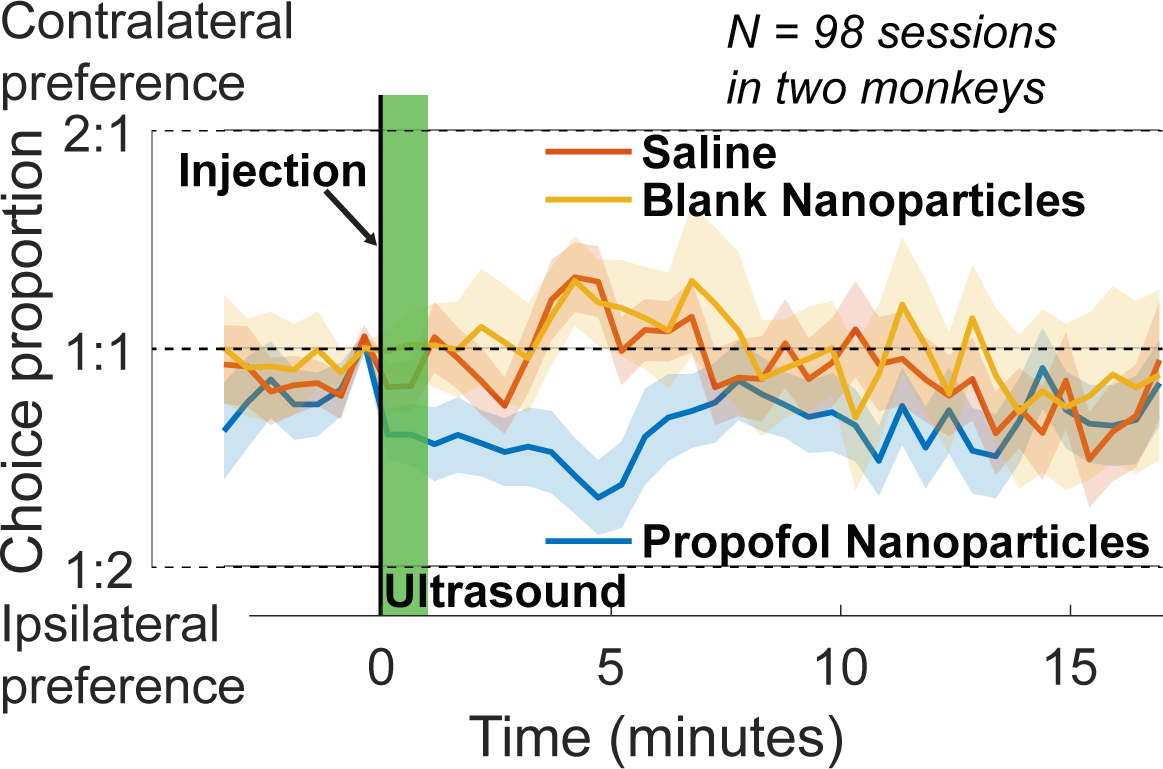
Propofol loaded in the nanoparticles is required for behavior modulation. Mean*±*s.e.m. proportion of choices contralateral to the targeted LGN as a function of time for the propofol-loaded nanoparticles (blue), saline (red), and blank nanoparticles (yellow). This plot incorporates all the data from Fig. 2 (1.2 and 1.5 MPa combined) as well as 18 sessions with blank nanoparticles (1.5 MPa only).

### Behavior Baseline Correction

The results shown in Fig. 2 contain sessions with baseline behavior periods which were highly biased. To assess the impact of these sessions, we have also analyzed the results by excluding any sessions in which the monkeys were biased more than 70% in either direction. Excluding these sessions removes the offset in baselines shown in Fig. 3, while retaining a clear bias induced by propofol delivery at the 1.5 MPa pressure. A two-way ANOVA with factors of LGN sonicated and injection type (the same analysis as the main text) detected a significant effect at 1.5 MPa 2-5 minutes after sonication (F(1,29) = 4.31, p = 0.047) but not 3-6 minutes after sonication (F(1,29) = 1.92, p = 0.18). However, the effect was not significant at 1.2 MPa with this analysis 2-5 minutes after sonication (F(1,29) = 3.52, p = 0.071). This suggests that for immediate effects to be substantial enough to modulate behavior, a pressure higher than 1.2 MPa may be necessary.

**Figure S5.**
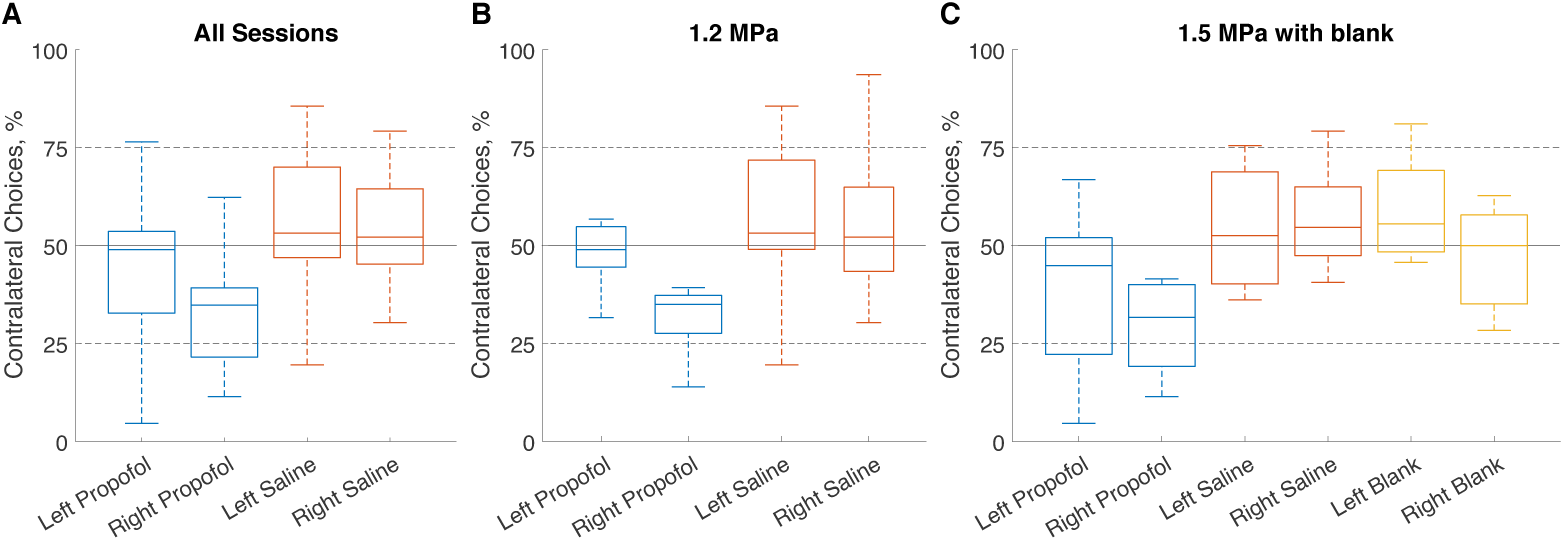
Statistical analysis at 5 minutes post-sonication used to quantify behavior bias. Box plots incorporating behavior bias in the time period 2-5 minutes after sonication for A) all sessions, B) 1.2 MPa sessions, and C) 1.5 MPa sessions. A two-factor ANOVA of side sonicated and type of injection was used to quantify the effects under these conditions.

**Figure S6.**
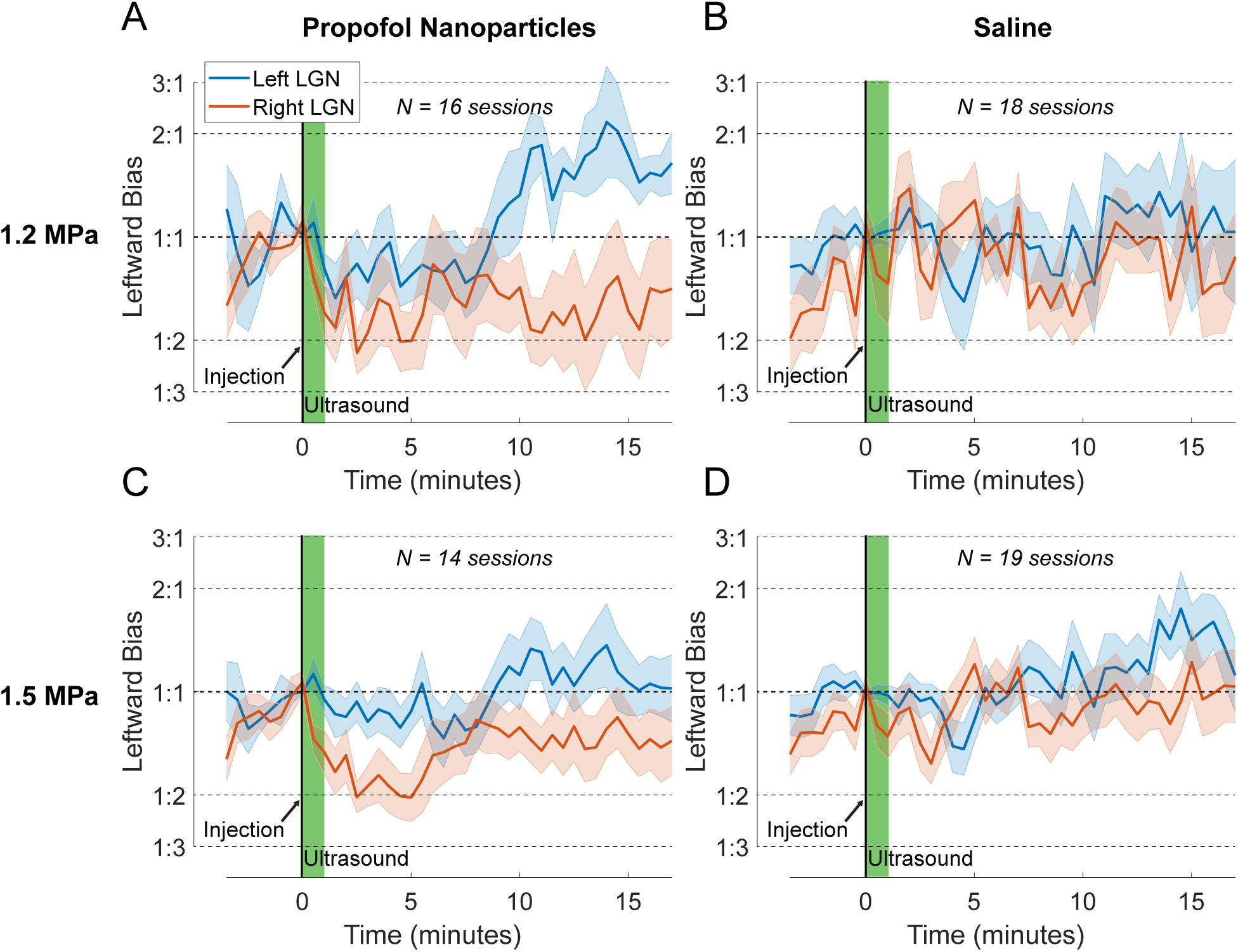
Propofol-induced modulation of choice behavior is target-specific also during sessions with a stable baseline. Same format as in Fig. 3, specifically using data that showed stable baseline, i.e., only including sessions with baseline bias less than 70% in either direction. The post-ultrasound effect is preserved and there is no tendency for a pre-ultrasound effect.

**Figure S7.**
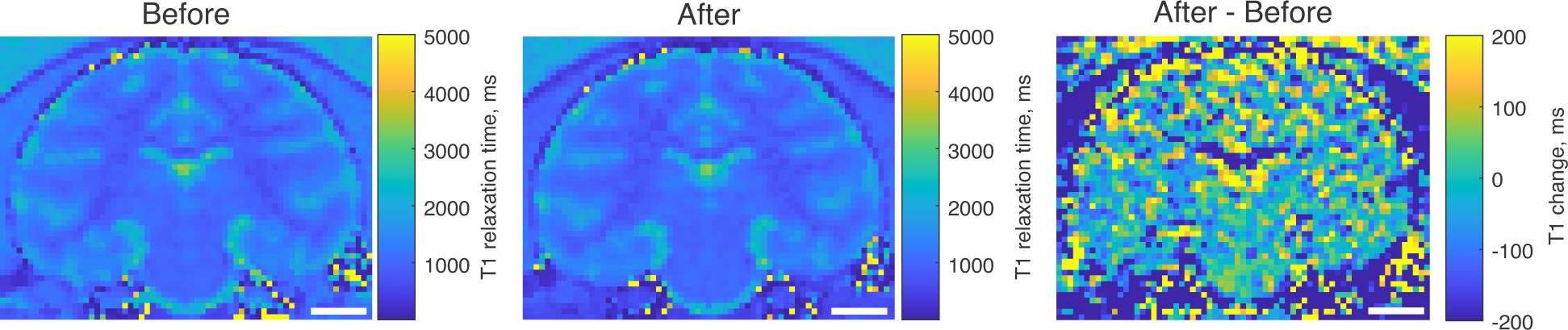
Quantitative T1 mapping shows no sign of BBB disruption in the sonicated LGN in Monkey 1. T1 relaxation time maps before (left), 30 minutes after ultrasound-triggered drug release and gadoteridol contrast administration (middle), and the difference between the two maps (right). The presence of gadolinium-based contrast would be expected to decrease T1 relaxation times in any regions of BBB disruption. Ultrasound was delivered to the left LGN.

**Figure S8.**
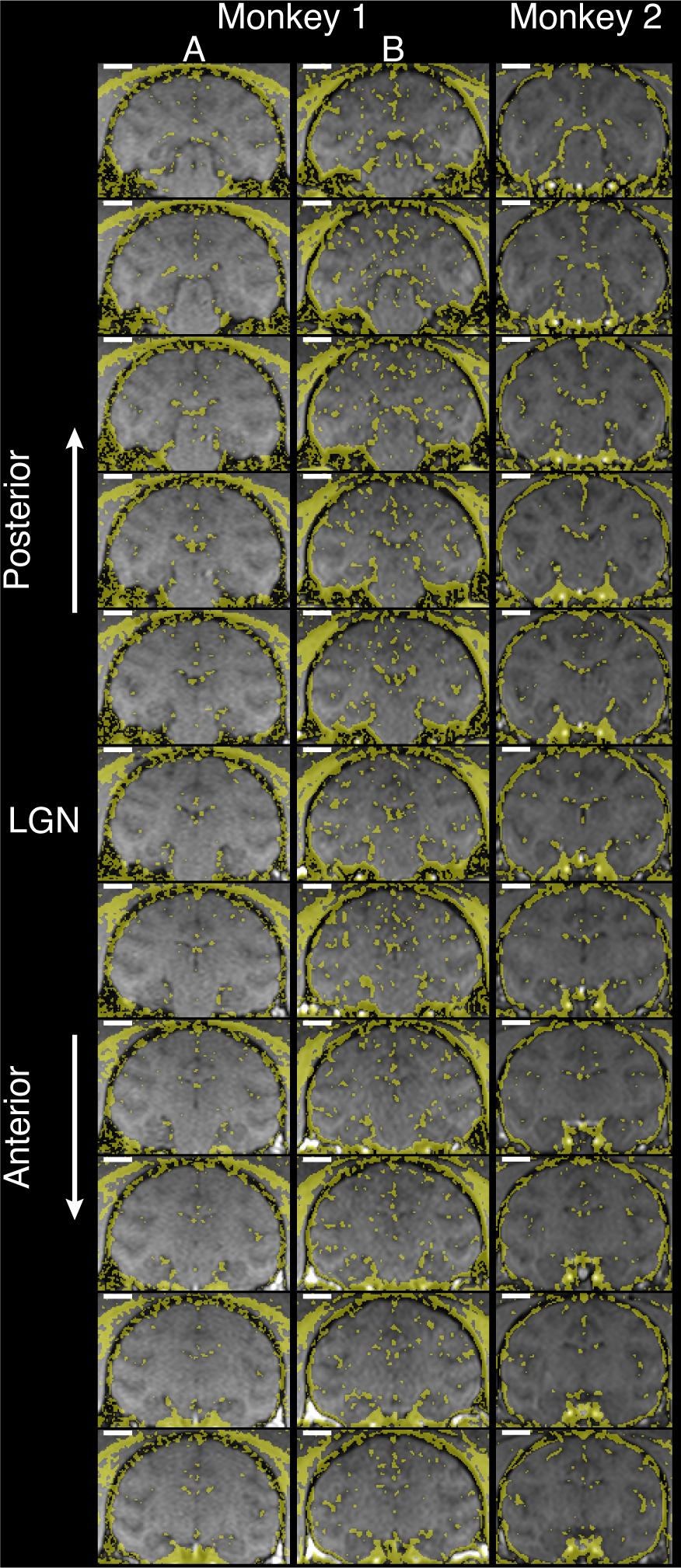
Gadolinium-enhanced T1-weighted MRI detected no sign of blood brain barrier disruption at the sonication target. Same analysis as in Fig. 4. MRI slices centered on the LGN (middle) and moving in 1 mm steps anteriorly (top) or posteriorly (bottom). The data are provided separately for the two monkeys and sessions A and B in each monkey (columns).

**Figure S9.**
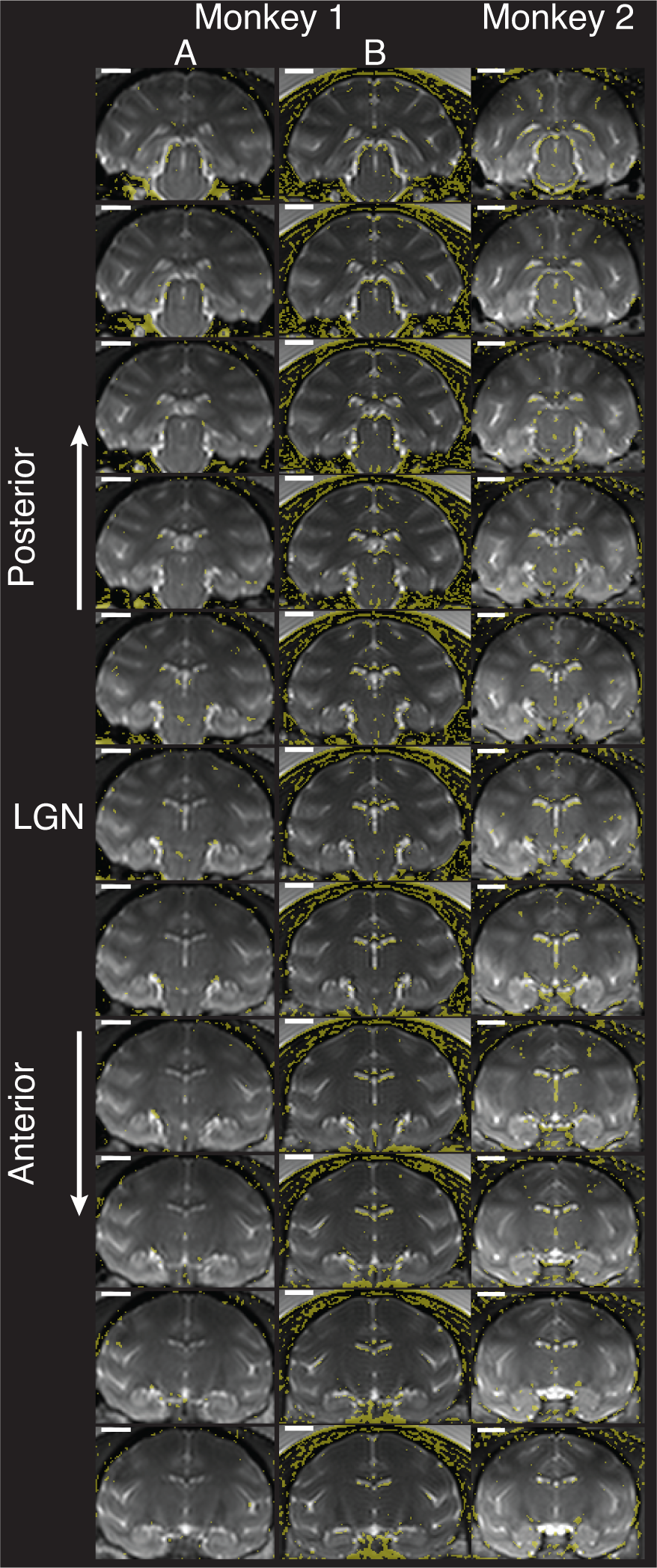
Gadolinium-enhanced T2-weighted MRI detected no sign of edema at the sonication target. Same format as in Fig. S8, using T2-weighted MRI.

**Table S1.**
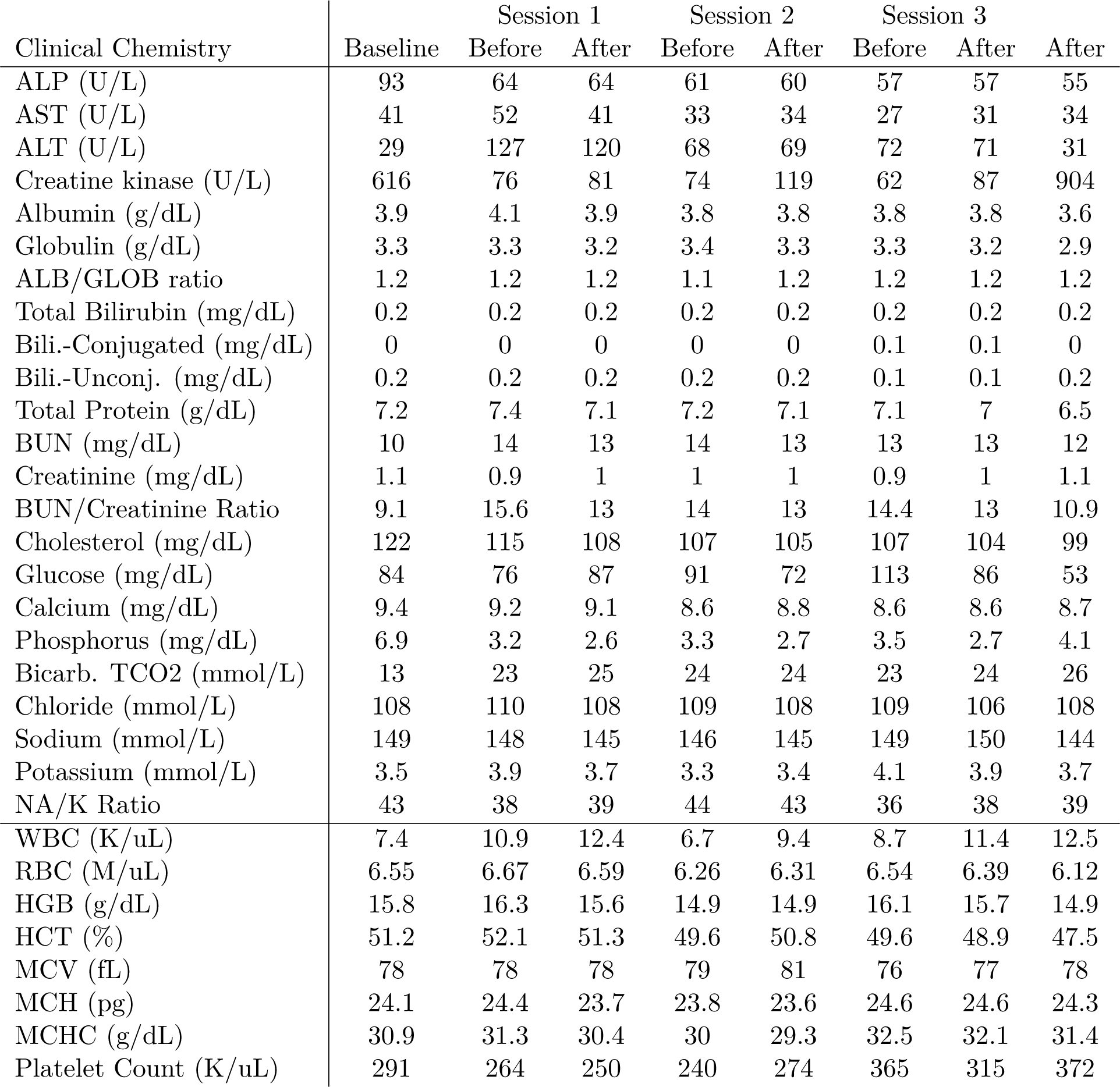
Clinical chemistry and hematology; Monkey 1. Clinical chemistry and hematology test outcomes based on blood draws taken during baseline, immediately before the injection of the PFOB-based nanoparticles, and 1.5 hours following the injection (columns). The data are provided separetely for the 3 sessions (columns). The blood was analyzed and values provided by IDEXX (Westbrook, Maine).

**Table S2.**
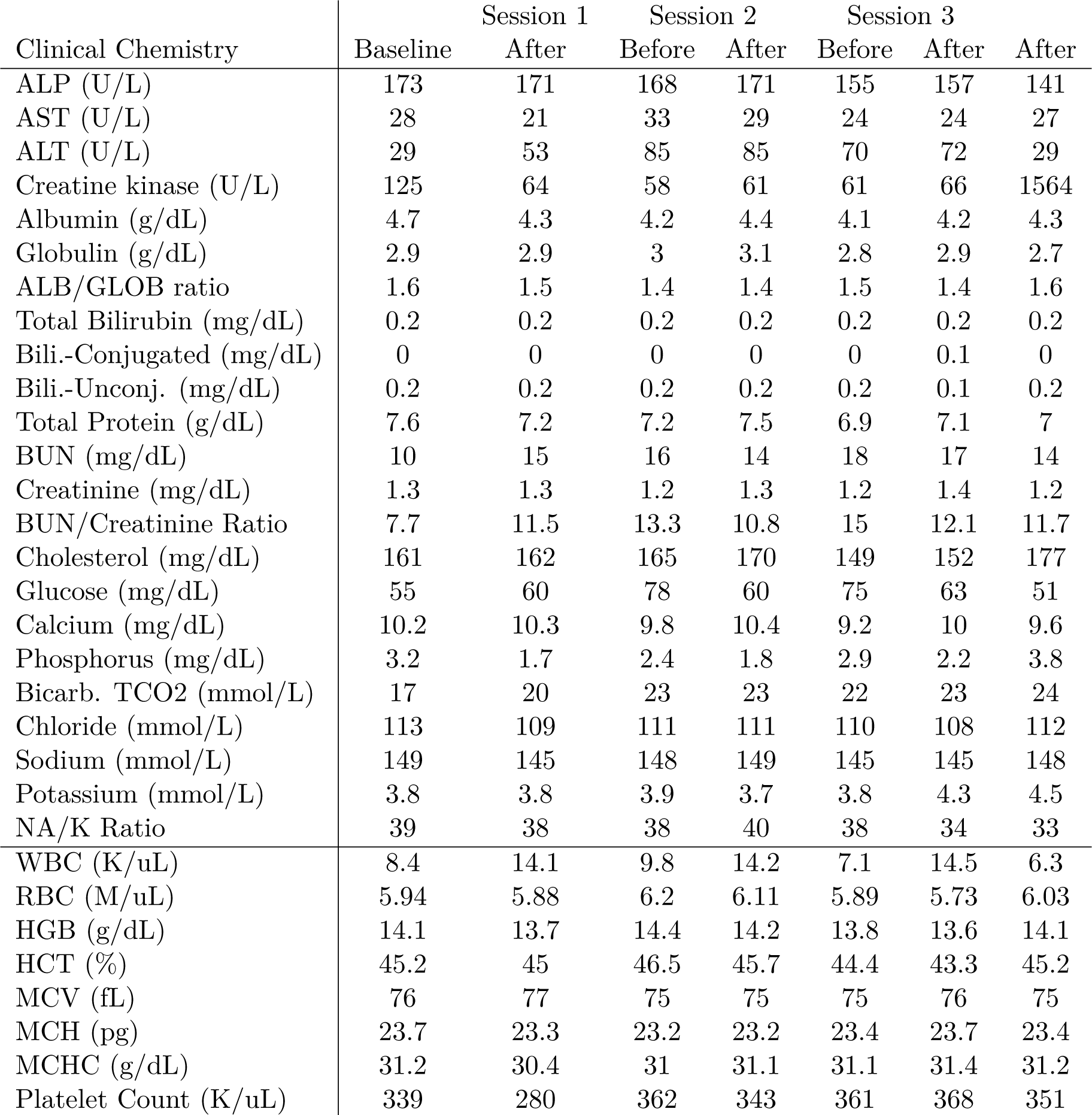
Clinical chemistry and hematology; Monkey 2. Same format as in Table S1 for Monkey 2.

## References

[1] Natalya Rapoport. Drug-loaded perfluorocarbon nanodroplets for ultrasound-mediated drug delivery. In Therapeutic Ultrasound, pages 221–241. Springer, 2016.

[2] Pejman Ghanouni, Kim Butts Pauly, W Jeff Elias, Jaimie Henderson, Jason Sheehan, Stephen Monteith, and Max Wintermark. Transcranial MRI-guided focused ultrasound: a review of the technologic and neurologic applications. American Journal of Roentgenology, 205(1):150–159, 2015.

[3] Oliver D Kripfgans, J Brian Fowlkes, Douglas L Miller, O Petter Eldevik, and Paul L Carson. Acoustic droplet vaporization for therapeutic and diagnostic applications. Ultrasound in medicine & biology, 26(7):1177–1189, 2000.

[4] Natalya Rapoport. Phase-shift, stimuli-responsive perfluorocarbon nanodroplets for drug delivery to cancer. Wiley Interdisciplinary Reviews: Nanomedicine and Nanobiotechnology, 4(5):492–510, 2012.

[5] Natalya Rapoport, Kweon-Ho Nam, Roohi Gupta, Zhongao Gao, Praveena Mohan, Allison Payne, Nick Todd, Xin Liu, Taeho Kim, Jill Shea, et al. Ultrasound-mediated tumor imaging and nanotherapy using drug loaded, block copolymer stabilized perfluorocarbon nanoemulsions. Journal of Controlled Release, 153(1):4–15, 2011.

[6] Cherry C Chen, Paul S Sheeran, Shih-Ying Wu, Oluyemi O Olumolade, Paul A Dayton, and Elisa E Konofagou. Targeted drug delivery with focused ultrasound-induced blood-brain barrier opening using acoustically-activated nanodroplets. Journal of Controlled Release, 172(3):795–804, 2013.

[7] Raag D Airan, Randall A Meyer, Nicholas PK Ellens, Kelly R Rhodes, Keyvan Farahani, Martin G Pomper, Shilpa D Kadam, and Jordan J Green. Noninvasive targeted transcranial neuromodulation via focused ultrasound gated drug release from nanoemulsions. Nano letters, 17(2):652–659, 2017.

[8] Jeffrey B Wang, Muna Aryal, Qian Zhong, Daivik B Vyas, and Raag D Airan. Noninvasive ultrasonic drug uncaging maps whole-brain functional networks. Neuron, 100(3):728–738, 2018.

[9] Harriet Lea-Banks, Meaghan A O’Reilly, Clement Hamani, and Kullervo Hynynen. Localized anesthesia of a specific brain region using ultrasound-responsive barbiturate nanodroplets. Theranostics, 10(6):2849, 2020.

[10] Harriet Lea-Banks, Ying Meng, Sheng-Kai Wu, Rania Belhadjhamida, Clement Hamani, and Kullervo Hynynen. Ultrasound-sensitive nanodroplets achieve targeted neuromodulation. Journal of Controlled Release, 332:30–39, 2021.

[11] Harriet Lea-Banks and Kullervo Hynynen. Sub-millimetre precision of drug delivery in the brain from ultrasound-triggered nanodroplets. Journal of Controlled Release, 338:731–741, 2021.

[12] Natalya Rapoport, Zhonggao Gao, and Anne Kennedy. Multifunctional nanoparticles for combining ultrasonic tumor imaging and targeted chemotherapy. Journal of the National Cancer Institute, 99(14):1095–1106, 2007.

[13] Paul S Larson. Deep brain stimulation for movement disorders. Neurotherapeutics, 11(3):465–474, 2014.

[14] Joseph L Price and Wayne C Drevets. Neural circuits underlying the pathophysiology of mood disorders. Trends in cognitive sciences, 16(1):61–71, 2012.

[15] Alik S Widge and Darin D Dougherty. Deep brain stimulation for treatment-refractory mood and obsessive-compulsive disorders. Current Behavioral Neuroscience Reports, 2(4):187–197, 2015.

[16] Urs Braun, Axel Schaefer, Richard F Betzel, Heike Tost, Andreas Meyer-Lindenberg, and Danielle S Bassett. From maps to multi-dimensional network mechanisms of mental disorders. Neuron, 97(1):14–31, 2018.

[17] Jens Kuhn, Wolfgang Gaebel, Joachim Klosterkoetter, and Christiane Woopen. Deep brain stimulation as a new therapeutic approach in therapy-resistant mental disorders: ethical aspects of investigational treatment. European Archives of Psychiatry and Clinical Neuroscience, 259(2):135–141, 2009.

[18] MP Dandekar, AJ Fenoy, AF Carvalho, JC Soares, and J Quevedo. Deep brain stimulation for treatment-resistant depression: an integrative review of preclinical and clinical findings and translational implications. Molecular psychiatry, 23(5):1094–1112, 2018.

[19] Katherine W Scangos, Ghassan S Makhoul, Leo P Sugrue, Edward F Chang, and Andrew D Krystal. State-dependent responses to intracranial brain stimulation in a patient with depression. Nature Medicine, 27(2):229–231, 2021.

[20] C Vanlersberghe and Frederic Camu. Propofol. Modern Anesthetics, pages 227–252, 2008.

[21] Marko M Sahinovic, Michel MRF Struys, and Anthony R Absalom. Clinical pharmacokinetics and pharmacodynamics of propofol. Clinical pharmacokinetics, 57(12):1539–1558, 2018.

[22] Claudia S Cohn and Melissa M Cushing. Oxygen therapeutics: perfluorocarbons and blood substitute safety. Critical care clinics, 25(2):399–414, 2009.

[23] Camila Irene Castro and Juan Carlos Briceno. Perfluorocarbon-based oxygen carriers: review of products and trials. Artificial organs, 34(8):622–634, 2010.

[24] Xiao Li, Zhongguo Sui, Xin Li, Wen Xu, Qie Guo, Jialin Sun, and Fanbo Jing. Perfluorooctylbromide nanoparticles for ultrasound imaging and drug delivery. International journal of nanomedicine, 13:3053, 2018.

[25] BA Orser, LY Wang, PS Pennefather, and JF MacDonald. Propofol modulates activation and desensitization of gabaa receptors in cultured murine hippocampal neurons. Journal of Neuroscience, 14(12):7747–7760, 1994.

[26] Qian Zhong, Byung C Yoon, M Aryal, Jeffrey B Wang, T Ilovitsh, MA Baikoghli, N Hosseini-Nassab, A Karthik, RH Cheng, KW Ferrara, et al. Polymeric perfluorocarbon nanoemulsions are ultrasound-activated wireless drug infusion catheters. Biomaterials, 206:73–86, 2019.

[27] Taylor D Webb, Matthew G Wilson, Henrik Odéen, and Jan Kubanek. Remus: System for remote deep brain interventions. Iscience, 25(11):105251, 2022.

[28] Taylor D Webb, Matthew G Wilson, Henrik Odeen, and Jan Kubanek. Sustained modulation of primate deep brain circuits with focused ultrasonic waves. Brain Stimulation, 2023.

[29] H Oppenheim. Über eine durch eine klinisch bisher nicht verwerthete Untersuchungsmethode ermittelte Form der Sensibilitätsstörung bei einseitigen Erkrankugen des Großhirns. Neurologisches Centralblatt, 4:529–533, 1885.

[30] Tony Ro, Chris Rorden, Jon Driver, and Robert Rafal. Ipsilesional biases in saccades but not perception after lesions of the human inferior parietal lobule. Journal of Cognitive Neuroscience, 13(7):920–929, 2001.

[31] Peter H Schiller and Edward J Tehovnik. Cortical inhibitory circuits in eye-movement generation. European Journal of Neuroscience, 18(11):3127–3133, 2003.

[32] Claire Wardak, Etienne Olivier, and Jean-René Duhamel. A deficit in covert attention after parietal cortex inactivation in the monkey. Neuron, 42(3):501–508, 2004.

[33] Jan Kubanek, Jingfeng M Li, and Lawrence H Snyder. Motor role of parietal cortex in a monkey model of hemispatial neglect. Proceedings of the National Academy of Sciences, 112(16):E2067–E2072, 2015.

[34] Laura D Lewis, Veronica S Weiner, Eran A Mukamel, Jacob A Donoghue, Emad N Eskandar, Joseph R Madsen, William S Anderson, Leigh R Hochberg, Sydney S Cash, Emery N Brown, et al. Rapid fragmentation of neuronal networks at the onset of propofol-induced unconsciousness. Proceedings of the National Academy of Sciences, 109(49):E3377–E3386, 2012.

[35] Fabrice Marquet, Tobias Teichert, Shih-Ying Wu, Yao-Sheng Tung, Matthew Downs, Shutao Wang, Cherry Chen, Vincent Ferrera, and Elisa E Konofagou. Real-time, transcranial monitoring of safe blood-brain barrier opening in non-human primates. PloS one, 9(2):e84310, 2014.

[36] Matthew E Downs, Amanda Buch, Carlos Sierra, Maria Eleni Karakatsani, Shangshang Chen, Elisa E Konofagou, and Vincent P Ferrera. Long-term safety of repeated blood-brain barrier opening via focused ultrasound with microbubbles in non-human primates performing a cognitive task. PloS one, 10(5):e0125911, 2015.

[37] Alison S Smith, Meredith A Weinstein, Michael T Modic, William Pavlicek, Lisa R Rogers, Thomas G Budd, Ronald M Bukowski, Joseph D Purvis, James K Weick, and Paul M Duchesneau. Magnetic resonance with marked t2-weighted images: improved demonstration of brain lesions, tumor, and edema. American journal of neuroradiology, 6(5):691–697, 1985.

[38] James J Choi, Mathieu Pernot, Scott A Small, and Elisa E Konofagou. Noninvasive, transcranial and localized opening of the blood-brain barrier using focused ultrasound in mice. Ultrasound in medicine & biology, 33(1):95–104, 2007.

[39] Ko-Ting Chen, Wen-Yen Chai, Ya-Jui Lin, Chia-Jung Lin, Pin-Yuan Chen, Hong-Chieh Tsai, Chiung-Yin Huang, John S Kuo, Hao-Li Liu, and Kuo-Chen Wei. Neuronavigation-guided focused ultrasound for transcranial blood-brain barrier opening and immunostimulation in brain tumors. Science Advances, 7(6):eabd0772, 2021.

[40] Yoshito Tsushima, Jun Aoki, and Keigo Endo. Brain microhemorrhages detected on t2*-weighted gradient-echo mr images. American journal of neuroradiology, 24(1):88–96, 2003.

[41] Yang Cao, Yuli Chen, Tao Yu, Yuan Guo, Fengqiu Liu, Yuanzhi Yao, Pan Li, Dong Wang, Zhigang Wang, Yu Chen, et al. Drug release from phase-changeable nanodroplets triggered by low-intensity focused ultrasound. Theranostics, 8(5):1327, 2018.

[42] Hak Soo Choi, Khaled Nasr, Sergey Alyabyev, Dina Feith, Jeong Heon Lee, Soon Hee Kim, Yoshitomo Ashitate, Hoon Hyun, Gabor Patonay, Lucjan Strekowski, et al. Synthesis and in vivo fate of zwitterionic near-infrared fluorophores. Angewandte Chemie International Edition, 50(28):6258–6263, 2011.

[43] George Paxinos, Xu-Feng Huang, and Arthur W Toga. The rhesus monkey brain in stereotaxic coordinates. 2000.

[44] Lauren D Martin, Gregory A Dissen, Matthew J McPike, and Ansgar M Brambrink. Effects of anesthesia with isoflurane, ketamine, or propofol on physiologic parameters in neonatal rhesus macaques (macaca mulatta). Journal of the American Association for Laboratory Animal Science, 53(3):290–300, 2014.

[45] Muna Aryal, Costas D Arvanitis, Phillip M Alexander, and Nathan McDannold. Ultrasound-mediated blood–brain barrier disruption for targeted drug delivery in the central nervous system. Advanced drug delivery reviews, 72:94–109, 2014.

[46] Nir Lipsman, Ying Meng, Allison J Bethune, Yuexi Huang, Benjamin Lam, Mario Masellis, Nathan Herrmann, Chinthaka Heyn, Isabelle Aubert, Alexandre Boutet, et al. Blood–brain barrier opening in alzheimer’s disease using mr-guided focused ultrasound. Nature communications, 9(1):2336, 2018.

[47] Maria Eleni Karakatsani, Javier Blesa, and Elisa Evgenia Konofagou. Blood–brain barrier opening with focused ultrasound in experimental models of parkinson’s disease. Movement Disorders, 34(9):1252–1261, 2019.

[48] Marie Pierre Krafft. Fluorocarbons and fluorinated amphiphiles in drug delivery and biomedical research. Advanced drug delivery reviews, 47(2-3):209–228, 2001.

[49] Phillip T Leese, Robert J Noveck, Jolene S Shorr, Catherine M Woods, Kathryn E Flaim, and Peter E Keipert. Randomized safety studies of intravenous perflubron emulsion. i. effects on coagulation function in healthy volunteers. Anesthesia & Analgesia, 91(4):804– 811, 2000.

[50] Robert J Noveck, EJ Shannon, Phillip T Leese, Jolene S Shorr, Kathryn E Flaim, Peter E Keipert, and Catherine M Woods. Randomized safety studies of intravenous perflubron emulsion. ii. effects on immune function in healthy volunteers. Anesthesia & Analgesia, 91(4):812–822, 2000.

[51] Shady Farah, Daniel G Anderson, and Robert Langer. Physical and mechanical properties of pla, and their functions in widespread applications—a comprehensive review. Advanced drug delivery reviews, 107:367–392, 2016.

[52] Vincent DeStefano, Salaar Khan, and Alonzo Tabada. Applications of pla in modern medicine. Engineered Regeneration, 1:76–87, 2020.

[53] Ruxandra Gref, Yoshiharu Minamitake, Maria Teresa Peracchia, Vladimir Trubetskoy, Vladimir Torchilin, and Robert Langer. Biodegradable long-circulating polymeric nanospheres. Science, 263(5153):1600–1603, 1994.

[54] Katrin Knop, Richard Hoogenboom, Dagmar Fischer, and Ulrich S Schubert. Poly (ethylene glycol) in drug delivery: pros and cons as well as potential alternatives. Angewandte chemie international edition, 49(36):6288–6308, 2010.

[55] Chetna Dhand, Molamma P Prabhakaran, Roger W Beuerman, R Lakshminarayanan, Neeraj Dwivedi, and Seeram Ramakrishna. Role of size of drug delivery carriers for pulmonary and intravenous administration with emphasis on cancer therapeutics and lung-targeted drug delivery. RSC advances, 4(62):32673–32689, 2014.

[56] Mazyar Malakouti, Archish Kataria, Sayed K. Ali, and Steven Schenker. Elevated Liver Enzymes in Asymptomatic Patients – What Should I Do? Journal of Clinical and Translational Hepatology, 5(4):394–403, December 2017. Publisher: Xia & He Publishing Inc.

[57] Richard M. Green and Steven Flamm. AGA technical review on the evaluation of liver chemistry tests. Gastroenterology, 123(4):1367–1384, October 2002.

[58] George Aragon and Zobair M. Younossi. When and how to evaluate mildly elevated liver enzymes in apparently healthy patients. Cleveland Clinic Journal of Medicine, 77(3):195–204, March 2010. Publisher: Cleveland Clinic Journal of Medicine Section: Review.

[59] Christoph Jacoby, Sebastian Temme, Friederike Mayenfels, Nicole Benoit, Marie Pierre Krafft, Rolf Schubert, Jürgen Schrader, and Ulrich Flögel. Probing different perfluoro-carbons for in vivo inflammation imaging by 19f mri: image reconstruction, biological half-lives and sensitivity. NMR in Biomedicine, 27(3):261–271, 2014.

[60] Lucie Somaglino, L Mousnier, A Giron, W Urbach, N Tsapis, and Nicolas Taulier. In vitro evaluation of polymeric nanoparticles with a fluorine core for drug delivery triggered by focused ultrasound. Colloids and Surfaces B: Biointerfaces, 200:111561, 2021.

[61] Paul S Sheeran, Samantha H Luois, Lee B Mullin, Terry O Matsunaga, and Paul A Dayton. Design of ultrasonically-activatable nanoparticles using low boiling point perfluorocarbons. Biomaterials, 33(11):3262–3269, 2012.

[62] Shih-Ying Wu, Samantha M Fix, Christopher B Arena, Cherry C Chen, Wenlan Zheng, Oluyemi O Olumolade, Virginie Papadopoulou, Anthony Novell, Paul A Dayton, and Elisa E Konofagou. Focused ultrasound-facilitated brain drug delivery using optimized nanodroplets: vaporization efficiency dictates large molecular delivery. Physics in Medicine & Biology, 63(3):035002, 2018.

[63] Jon N Marsh, Christopher S Hall, Samuel A Wickline, and Gregory M Lanza. Temperature dependence of acoustic impedance for specific fluorocarbon liquids. The Journal of the Acoustical Society of America, 112(6):2858–2862, 2002.

[64] Pelle Ohlsson, Klara Petersson, Per Augustsson, and Thomas Laurell. Acoustic impedance matched buffers enable separation of bacteria from blood cells at high cell concentrations. Scientific reports, 8(1):9156, 2018.

[65] Takeshi Kubota, Yuta Kurashina, JianYi Zhao, Keita Ando, and Hiroaki Onoe. Ultrasound-triggered on-demand drug delivery using hydrogel microbeads with release enhancer. Materials & Design, 203:109580, 2021.

[66] J Kanto and E Gepts. Pharmacokinetic implications for the clinical use of propofol. Clinical pharmacokinetics, 17:308–326, 1989.

[67] YS Tapia-Guerrero, Del Prado-Audelo, FV Borbolla-Jiménez, DM Giraldo Gomez, I García-Aguirre, CA Colín-Castro, JA Morales-González, G Leyva-Gómez, JJ Magaña, et al. Effect of uv and gamma irradiation sterilization processes in the properties of different polymeric nanoparticles for biomedical applications. Materials, 13(5):1090, 2020.

[68] US Food, Drug Administration, et al. Bacterial endotoxins/pyrogens. Inspections, Compliance, Enforcement, and Criminal Investigations, FDA, Bethesda, MD: http://www.fda.gov/ICECI/Inspections/InspectionGuides/InspectionTechnica, 1985.

[69] Jan Kubanek, Julian Brown, Patrick Ye, Kim Butts Pauly, Tirin Moore, and William Newsome. Remote, brain region–specific control of choice behavior with ultrasonic waves. Science Advances, 6(21):eaaz4193, 2020.

[70] Meaghan A O’Reilly, Aidan Muller, and Kullervo Hynynen. Ultrasound insertion loss of rat parietal bone appears to be proportional to animal mass at submegahertz frequencies. Ultrasound in medicine & biology, 37(11):1930–1937, 2011.

[71] Charlotte Constans, Thomas Deffieux, Pierre Pouget, Mickael Tanter, and Jean-François Aubry. A 200–1380-khz quadrifrequency focused ultrasound transducer for neurostimulation in rodents and primates: transcranial in vitro calibration and numerical study of the influence of skull cavity. IEEE transactions on ultrasonics, ferroelectrics, and frequency control, 64(4):717–724, 2017.

[72] Axel Montagne, Arthur W Toga, and Berislav V Zlokovic. Blood-brain barrier permeability and gadolinium: benefits and potential pitfalls in research. JAMA neurology, 73(1):13–14, 2016.

[73] Min-Chi Ku, Sonia Waiczies, Thoralf Niendorf, and Andreas Pohlmann. Assessment of blood brain barrier leakage with gadolinium-enhanced mri. Preclinical MRI: methods and protocols, pages 395–408, 2018.

[74] Thomas J Manuel, Michelle K Sigona, M Anthony Phipps, Jiro Kusunose, Huiwen Luo, Pai-Feng Yang, Allen T Newton, John C Gore, William Grissom, Li Min Chen, et al. Small volume blood-brain barrier opening in macaques with a 1 mhz ultrasound phased array. Journal of Controlled Release, 363:707–720, 2023.

[75] Carmen Gasca-Salas, Beatriz Fernández-Rodríguez, José A Pineda-Pardo, Rafael Rodríguez-Rojas, Ignacio Obeso, Frida Hernández-Fernández, Marta Del Álamo, David Mata, Pasqualina Guida, Carlos Ordás-Bandera, et al. Blood-brain barrier opening with focused ultrasound in parkinson’s disease dementia. Nature communications, 12(1):779, 2021.

[76] Graziella Donatelli, Paolo Cecchi, Gianmichele Migaleddu, Matteo Cencini, Paolo Frumento, Claudio D’Amelio, Luca Peretti, Guido Buonincontri, Livia Pasquali, Michela Tosetti, et al. Quantitative t1 mapping detects blood–brain barrier breakdown in apparently non-enhancing multiple sclerosis lesions. NeuroImage: Clinical, 40:103509, 2023.

[77] Rashi I Mehta, Jeffrey S Carpenter, Rupal I Mehta, Marc W Haut, Manish Ranjan, Umer Najib, Paul Lockman, Peng Wang, Pierre-François D’haese, and Ali R Rezai. Blood-brain barrier opening with mri-guided focused ultrasound elicits meningeal venous permeability in humans with early alzheimer disease. Radiology, 298(3):654–662, 2021.

